# Broad-spectrum *in situ* genetic reprogramming of native microbiomes

**DOI:** 10.64898/2026.11.03.764085

**Authors:** Jeongsoo Hur, Charles Sanfiorenzo, Benjamin Simpson, Anastasiya Oguienko, Jianyi Huang, Mikel Lipschitz, Raymond J. Zhang, Tomasz Gawda, Sarkis K. Mazmanian, Kaihang Wang

## Abstract

Complex microbial communities are found throughout terrestrial, aquatic, and animal-associated ecosystems. Genetic modification of discrete microbial taxa within native communities represents a powerful tool to study and augment microbiome functions. Herein, we develop Synthetic Phage Assisted Reprogrammable and Targetable Addition (SPARTA) as a one-step strategy for packaging, delivering, and integrating large synthetic DNA into a broad range of native microbiomes. SPARTA leverages a broad-spectrum synthetic phage particle for *in situ* genome engineering within complex soil, freshwater, and mammalian fecal microbiomes, reprogramming 59 native species across 16 families. By modulating SPARTA components, we further demonstrated tunability of SPARTA towards targeted microbiome population. Additionally, we built a novel companion tool to quantitatively identify engineered taxa, named “16S crRNA-guided inference of bacterial engineering sequencing” or ScRibe-seq. The SPARTA-ScRibe-seq pipeline delivered and validated functional synthetic gene clusters in diverse native microbiomes while preserving the original microbial community structure. Together, this novel genome engineering framework constitutes an end-to-end programmable platform for rational and customizable reprogramming of native microbiomes.

## Main

Microbial communities are ubiquitous across both abiotic and biotic environments, including soils^1–3^, aquatic systems^4,5^, and the tissues of fungi^6^, plants^7,8^, and animals^9,10^. Microbiome consortia impact their habitats, modulate host physiology, and ultimately shape the function and structure of entire ecosystems^11^. Given their high abundance in diverse environments (estimated at 10^5^–10^12^ bacteria per gram of environmental material^5,12^), microbial consortia can be modified for environmental^3,13^ and biomedical applications^14,15^ through genome rewriting. In this work, we aim to expand our capacity to engineer native microbiome within their native environments at the community scale.

Previous studies attempted bacterial conjugation to transfer synthetic DNA into laboratory strains or native bacteria^16–18^. In these approaches, donor bacteria conjugate DNA in nutritionally rich environments, such as the murine gut^16,18^. To address concerns of introducing such engineered donor bacteria in the native environment^19,20^, bacteriophages are being developed as an alternative delivery vehicles, harnessing lambda and T-series phages for antimicrobials development^21^ and for base editing^22^ in *E. coli*-colonized murine models. Despite the promise of this new paradigm, phage-based native microbiome engineering remains constrained by two major bottlenecks: 1) Achieving programmable delivery of phage over a broad host range; and 2) The limited genome-engineering capacities available to program new functions. Therefore, the ability to implement and analyze phage-mediated genetic engineering across diverse native microbiome communities is still limited.

To overcome these bottlenecks, we propose the Synthetic Phage Assisted Reprogrammable and Targetable Addition (SPARTA) system. By utilizing a P1 phage-like particle with broader tropism and high delivery capacity (∼100-kb)^23,24^, SPARTA is capable of delivering large DNA cargos across a wide range of microbiome taxa. In addition, a CRISPR-associated transposase (CAST)^25–27^ is adopted as a targetable genome engineering tool due to its broad compatibility and scalable integration of cargo. This unique combination synergistically allows SPARTA to achieve robust *in situ* reprogramming of genetic elements within diverse environment-associated microbiome consortia with tunable specificity and scalable capacity (**Fig. 1a**, Designing SPARTA).

**Figure 1.**
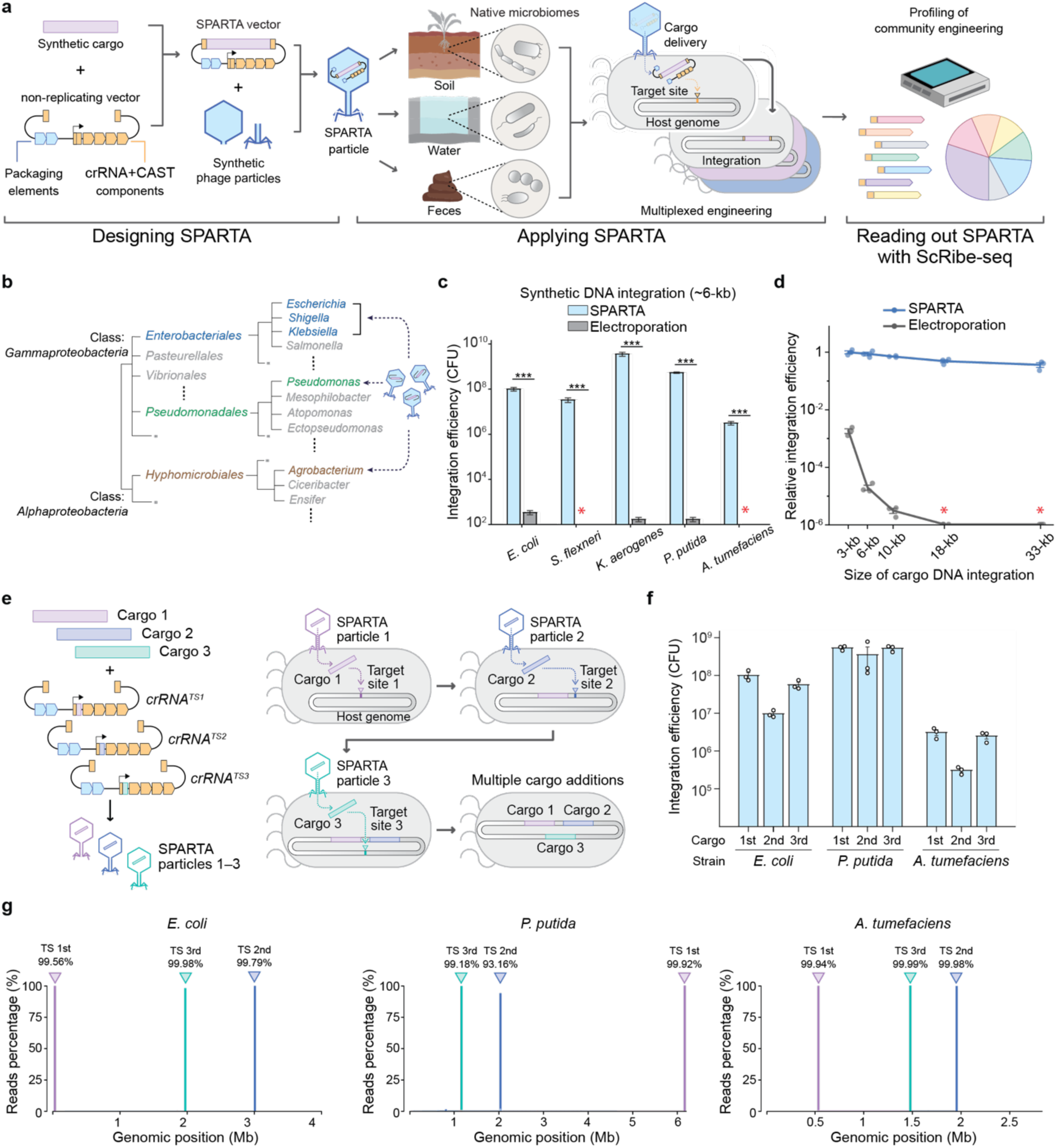
Workflow of the SPARTA system for *in situ* microbiome engineering and *in vitro* validation. **(a)** Summarized workflow of *in situ* multiplexed genome engineering with SPARTA from design, application to readout with ScRibe-seq. An “all-in-one”, non-replicating SPARTA vector contains cargo DNA, CRISPR-transposon (CAST) genome integration machinery, a crRNA programming the target site (TS), and phage packaging elements. SPARTA particles are applied to diverse microbiome samples *in situ*, donating cargo DNA flanked by two transposon ends into a target locus. SPARTA engineering is analyzed with ScRibe-seq to quantitatively profile engineered microbiome taxa at high resolution. **(b)** Phylogenetic tree showing the relationship between *Gammaproteobacteria* and *Alphaproteobacteria* classes, highlighting species applied with SPARTA particles. **(c)** Integration efficiency of synthetic DNA (∼6-kb) into several *Enterobacteriales* bacteria cells using SPARTA particles and electroporation. Red asterisk represents an integration efficiency of zero. **(d)** Relative integration efficiency as a function of cargo DNA size resulting from integration experiments via SPARTA and electroporation. Red asterisk represents an integration efficiency of zero. **(e)** Schematic of iterative DNA integration workflow via SPARTA. SPARTA particles packaging individual cargo DNA and corresponding target sites were manufactured separately and applied sequentially to recipient cells. The cargos included T7 RNA polymerase, SP6 RNA polymerase, and the π replicator, common synthetic bacterial modules used in biotechnology. **(f)** Integration efficiency of sequential engineering rounds in *E. coli*, *P. putida*, and *A. tumefaciens*. **(g)** Tn-seq of iterative genome engineering of *E. coli*, *P. putida*, and *A. tumefaciens*. On-target integration efficiency of cargos into each target site across distinct genomic positions. All error bars indicate the mean ± standard error of the mean (SEM) from three experimental replicates (N = 3). Three asterisks indicate a P <0.001, based on the Student’s t-test or one-way ANOVA.

We applied SPARTA across a wide variety of habitats, including soil, water, and feces to demonstrate its versatility and reliability (**Fig. 1a**, Applying SPARTA). Using SPARTA, we achieved multiplexed *in situ* genome engineering, resulting in modification of 59 diverse species spanning 16 different families within native microbiomes without disturbing taxonomic composition. This strategy of native consortia engineering via SPARTA enables engineering events to be readily detectable without selection or enrichment, allowing for the unbiased verification and interpretation of microbiome modification.

A central challenge for *in situ* microbial community engineering is the lack of a systematic strategy to read out the performance of metagenomic modification^17,28^. Quantitative profiling of the engineering outcome within native and complex microbial populations is essential to evaluate design and operation efficacy. Conventional readouts rely on antibiotic selection or culture-based assays, introducing strong biases and failing to capture community-level dynamics^29^. To address this, we developed “16S crRNA-guided inference of bacterial engineering sequencing” (ScRibe-seq) to comprehensively read out SPARTA engineering events across community members in native contexts (**Fig. 1a**, Reading out SPARTA with ScRibe-seq). By specifically amplifying DNA integration signature and 16S rRNA gene in tandem through ScRibe-seq, multiplexed engineering of microbiome communities can be profiled at high resolution.

Designing, applying, and reading out SPARTA events encompass a unique full-stack pipeline that enables rationally customizable reprogramming of native environment and host-associated microbiomes at the community scale. This new paradigm ushers in novel possibilities for environmental and clinical applications by unlocking our ability to write designed functions in native bacteria communities.

## Results

### Design of the SPARTA system for streamlined *in situ* engineering

The SPARTA system installs synthetic cargo using a packaging and delivery component derived from P1 phage, and a genome integration component utilizing CAST (**Fig. 1a**). We chose the P1 phage structure as a starting point to generate synthetic phage particles carrying the SPARTA vector, herein SPARTA particles, due to its broad tropism and large capacity (∼100-kb). Unlike phages interacting with specific host receptors (*e.g., LamB* for lambda^30^ and *OmpC* for T4^31^ phages), SPARTA particles recognize widespread sugar motifs in core lipopolysaccharide (LPS)^32,33^, which broadens the host range of engineering. In addition, a CAST component was employed for scalable and programmable large cargo integration, motivated by prior demonstrations of mega-base scale DNA integration across diverse prokaryotes^34^. To program complexed functions in native microbiomes, an “all-in-one” SPARTA vector was encapsulated by phage particles encoding both the functional DNA payload and CAST machinery (**Fig. 1a**, Designing SPARTA). After the SPARTA vector is delivered into recipient bacteria, the cargo DNA can be inserted in a target site of the genome, which is pre-defined by the CRISPR RNA (crRNA) of CAST.

These key components of SPARTA are further modified to be nonpersistent when applied to environments for *in situ* applications. SPARTA particles cannot spontaneously propagate due to the removal of the *coi* gene (P1 Δ*coi)*^35,36^, which is necessary to induce the lytic cycle of the phage, thereby reducing risks of downstream virulent phage contamination in treated environments. In addition, SPARTA vectors are modified to be non-replicating within native microbiomes in the absence of DNA integration, as the backbone is based on the *R6K* origin of replication, which cannot be maintained in hosts lacking the π replicator gene (*piR*). To manufacture these nonpersistent particles, we constructed a synthetic phage genome (P1 Δ*coi*::*piR pacA\**) that supplies *piR* proteins and encodes a mutated *pacA* variant^37^ with enhanced packaging efficiency (**Extended Data Fig. 1a**). Notably, the resulting non-replicating *R6K*-based SPARTA particles exhibit no observable compromise in DNA delivery efficiency compared to phage particles carrying replicating vectors (**Extended Data Fig. 1b**), while preventing false positives arising from the episomal maintenance of selection markers (**Extended Data Fig. 1c**). These feats substantially improve on the selectivity of cargo integration, while allowing SPARTA to remain minimally invasive for *in situ* engineering applications in native environments.

### *In vitro* characterization of SPARTA in different species, orders and classes

SPARTA is a broad-spectrum and efficient genome engineering system, capable of integrating large DNA cargos across different species, orders and classes. To explore this, we set out to test *in vitro* SPARTA engineering in distinct recipient targets spanning *Enterobacteriales* – the canonical P1 phage hosts *Escherichia coli*, *Shigella flexneri*, and *Klebsiella aerogenes* – and extending across orders to *Pseudomonas putida* (*Pseudomonadales*) and across classes to *Agrobacterium tumefaciens* (*Alphaproteobacteria*), two widely used chassis in soil microbiome studies^38,39^ (**Fig. 1b**). While electroporation often failed to yield integrants, likely due to lower delivery efficiencies, SPARTA achieved successful genome integration of synthetic cargo (∼6-kb) in all tested species with significantly higher integration efficiencies (**Fig. 1c** and **Extended Data Fig. 1d**). To our knowledge, this data provides the first direct evidence of P1-derived phage particle-mediated DNA integration into the *Alphaproteobacteria* class which is beyond the traditional P1 host range within the *Gammaproteobacteria* class.

Additionally, SPARTA can effectively introduce large genetic cassettes due to its payload scalability. To assess this, cargo DNAs ranging from 3 to 33-kb were integrated into the *E. coli* genome through SPARTA (**Fig. 1d** and **Extended Data Fig. 1e**), showcasing robust integration efficiencies independent of cargo size. Highly efficient integration of large DNA (∼33-kb) demonstrates SPARTA’s capability to install multi-gene pathways or other large genetic programs in a single step, owing to the combined high packaging capacity of P1-derived particles (up to 100-kb) and cargo tolerance of CAST^34^.

We further demonstrated that SPARTA is capable of iterative operation. Sets of SPARTA particles carrying distinct cargos (cargos 1–3) were programmed to target three non-overlapping genomic loci (target sites; TS 1–3) in each target recipient (**Fig. 1e**). Model microbiomes were sequentially treated with each SPARTA particle, and transposon-end sequencing (Tn-seq)^25^ was performed after every round of engineering to quantify on-target precision for each newly installed insertion. All microbes tested exhibited high integration efficiency (10^6^ – 10^8^ CFU per reaction) and on-target precision (93 – 99%) regardless of the cargo contents, size, target site, and round of insertion (**Figs. 1f–g**). Collectively taken, these results indicate that the SPARTA system enables robust *in vitro* engineering performance for large foreign DNA integrations in diverse bacterial cells.

### *In situ* SPARTA engineering of native microbiomes

SPARTA particles unlock the ability to engineer native microbiome communities by crRNA-mediated targeting of universally conserved sequences. Motivated by its flexibility and robustness *in vitro*, the SPARTA system was further designed to direct DNA cargo integration across complex microbiome consortia, in contrast to previous approaches that were limited to individual species or synthetic communities^17,22^. To achieve this, we identified universal target sites (UTS), such as highly conserved sequences presented within prokaryotic 16S–23S rRNA operons (**Fig. 2a**). Four UTS-specific crRNAs (UTS1–4) were individually designed and cloned into SPARTA vectors to enable multiplexed genome engineering of native bacterial communities.

**Figure 2.**
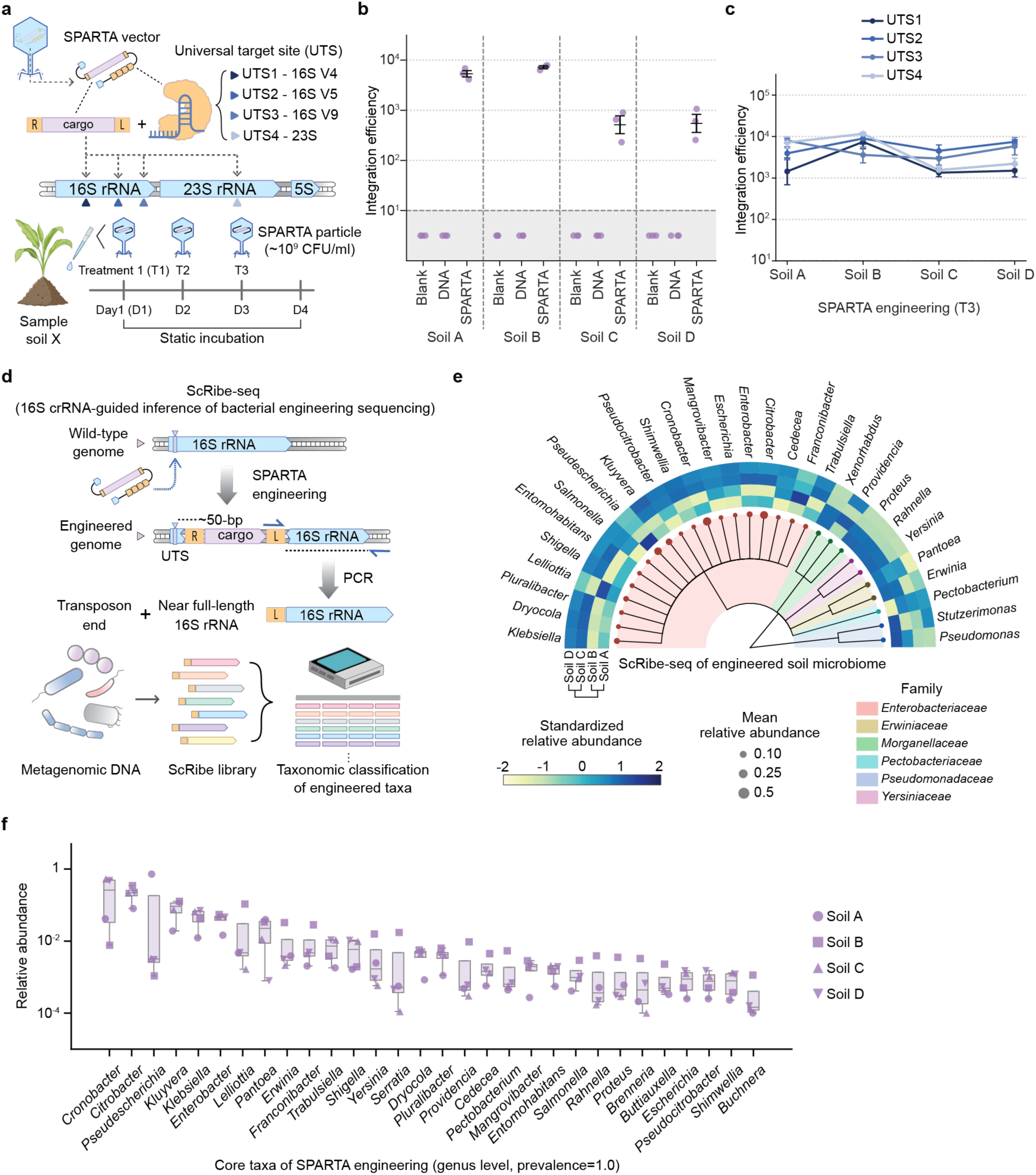
*In situ* microbiome community engineering with SPARTA and analysis of engineering profiles through ScRibe-seq. **(a)** Experimental scheme of multiplexed *in situ* engineering via SPARTA with crRNA targeting conserved sequences within the 16S-23S operon. Soil samples were treated with 10^9^ CFU/ml of SPARTA particles once a day and then collected for downstream analysis one day after the last SPARTA treatment. **(b)** SPARTA integration efficiency in all soil samples treated with the T3 dosage of blank solution, unpackaged DNA control, and SPARTA particles. The efficiency was calculated from the logarithm of CFU per gram of soil. All error bars indicate the mean ± standard error of the mean (SEM) from three experimental replicates (N = 3). **(c)** Integration efficiency of different soil samples engineered by SPARTA using various crRNA targets. All error bars indicate the mean ± SEM (N = 3). **(d)** Schematic of the ScRibe-seq workflow, enabling quantification and taxonomic profiling of prokaryotic species engineered by the SPARTA system. After SPARTA engineering *in situ*, the cargo flanked by two transposon ends is integrated approximately 50-bp downstream of the crRNA target site. ScRibe-seq employs one primer annealing to the transposon end flanking the inserted cargo DNA and another primer annealing to a highly conserved 16S rRNA sequence adjacent to the UTS, thereby selectively amplifying prokaryotic genomes containing SPARTA-programmed insertions. A near full-length of 16S rRNA (1,100–1,400-bp) with a CAST transposon end is selectively PCR-amplified from the engineered metagenomics. The resulting ScRibe library is analyzed with ONT Nanopore long-read sequencing for taxonomic classification. R; right transposon end, L; left transposon end. **(e)** 16S taxonomic classification of ScRibe-seq data at the genus level for different soil samples, displayed with a phylogenetic tree colored by family level. The standardized relative abundance of each taxon is shown as a heat map, and the size of the nodes represents the mean relative abundance of the taxa across samples. **(f)** Relative abundance of core genera engineered by SPARTA across all soil samples tested (N = 4). Boxes span from the 25th to 75th percentiles (interquartile range), whiskers extend to 1.5 times the interquartile range, and center lines represent the median.

Multiplexed engineering of microbiome communities in soil samples was achieved by using SPARTA particles targeting UTS. As an initial attempt, soil was selected as a representative environmental sample to evaluate SPARTA engineering capabilities *in situ*, given its high species diversity and the abundance of the *Pseudomonadota* phylum, which are putative targets of the current SPARTA design. To emulate field application conditions, SPARTA particle resuspensions were applied directly to soil once daily without agitation (**Fig. 2a**). Four different natural soil samples (soils A–D) collected from different geographical locations were treated with SPARTA particles under three dosing regimens (T1–T3, **Fig. 2a**). Different dosing regimens were carried out due to a potential for SPARTA iteration and an expected increase in SPARTA particle transduction events as a function of dosage. SPARTA engineering yielded consistent DNA integration into native microbiomes in all soil samples collected, whereas a blank solution and naked DNA (unpackaged SPARTA vector alone) exhibited no detectable integration events (**Fig. 2b**), confirming the genomic engineering was driven by SPARTA particles. We further demonstrated that the efficiency of engineering increases in a dose-dependent manner as a function of the number of treatments or phage particle titer (**Extended Data Figs. 2a–b**). Successful targeting of alternative UTS within the 16S–23S rRNA likewise supported robust and consistent multiplexed DNA integration across soil microbiome communities (**Fig. 2c**).

### ScRibe-seq enables taxonomic profiling and quantification of SPARTA engineering

SPARTA’s unique design enables profiling of engineered native species via ScRibe-seq. SPARTA engineers native bacterial communities without the use of living donors nor the uncontrolled spread of genetic cargo within environments, allowing us to develop the first high-throughput NGS approach capable of directly detecting targeted genome integration events in native contexts at the taxonomic level. The high-resolution profiling of genetically edited microbiomes across different samples and treatment conditions is essential for predictive control of multiplexed engineering outcomes^28^. Towards this end, ScRibe-seq was developed by coupling the concept of 16S rRNA gene amplicon sequencing with transposon end sequencing (**Fig. 2d)**. A ScRibe-seq library is generated by selectively amplifying the CAST transposon end adjacent to a near full-length sequence of the 16S rRNA gene target, which results from successful SPARTA engineering events in prokaryotic genomes. The resultant amplicon includes most hypervariable regions that enable robust taxonomic classification by using Oxford Nanopore long-read sequencing (**Fig. 2d**). We note that ScRibe-seq enables libraries to be generated directly from metagenomic DNA extracted from the *in situ* engineered samples without any enrichment, minimizing biases that conventionally result from laboratory culture expansion. The ScRibe-seq method thereby allows us to effectively analyze the *in situ* engineering profile of SPARTA at high resolution from complex, native microbial DNA mixtures.

By harnessing the ScRibe-seq strategy, the taxonomic footprint of *in situ* SPARTA engineering revealed variable patterns in the relative abundance of engineered taxa. Across all soil samples, a total of 49 genera spanning 9 families were identified as SPARTA-engineered microbes, demonstrating broad and multiplexed engineering of native bacterial community members (**Fig. 2e** and **Extended Data Fig. 2c**). Additionally, a core set of the 30 most abundant, consistently edited genera (prevalence = 1.0 across soil samples) showed that the majority of high-abundance taxa belonged to the *Enterobacteriaceae* family, such as *Citrobacter and Pseudescherichia* (**Fig. 2f**). This result is consistent with our current SPARTA design, in which both the phage particle and vector components are optimized for *Enterobacteriaceae*. Despite this design bias, ScRibe-seq also detected edited taxa outside *Enterobacteriaceae*, underscoring the capacity of SPARTA for broad-spectrum engineering in complex communities. Notably, the composition of edited taxa varied between soil samples, suggesting that community context, *e.g.,* local baseline relative abundance, modulates effective editing range. Together, these results demonstrate that SPARTA engineers genomes *in situ* across diverse taxa while generating soil-dependent, edited community profiles.

### Versatility of SPARTA for *in situ* engineering across diverse natural sample types

SPARTA is generalizable to a wide range of environmental contexts, each characterized by distinct microbial compositions. To assess this, we applied SPARTA to three natural sample types – extending beyond soil to further encompass natural freshwater and mammalian fecal samples (**Fig. 3a** and **Extended Data Figs. 3a–b**). All samples were subjected to static incubation with phage particles targeting UTS3 for community engineering. Principal coordinates analysis (PCoA) of ScRibe-seq profiling demonstrated robust engineering across all sample types, with engineered communities grouped by sample origin as shown in 16S beta diversity clustering (**Figs. 3b–c**; PERMANOVA, *P* = 0.0035, *R²* = 0.8283). In addition, SPARTA treatment in different environments showed negligible perturbation of the original microbial populations (**Extended Data Fig. 3c**), evidencing minimally invasive *in situ* engineering. Interestingly, centered log ratio (CLR) comparison and ALDEx2^40^ differential abundance screening further revealed several sample type-specific engineered taxa: *Pseudomonas* in soil; *Aeromonas, Alishewanella*, and *Rheinheimra* in water; and *Serratia* in fecal samples (**Fig. 3d** and **Extended Data Fig. 3d**).

**Figure 3.**
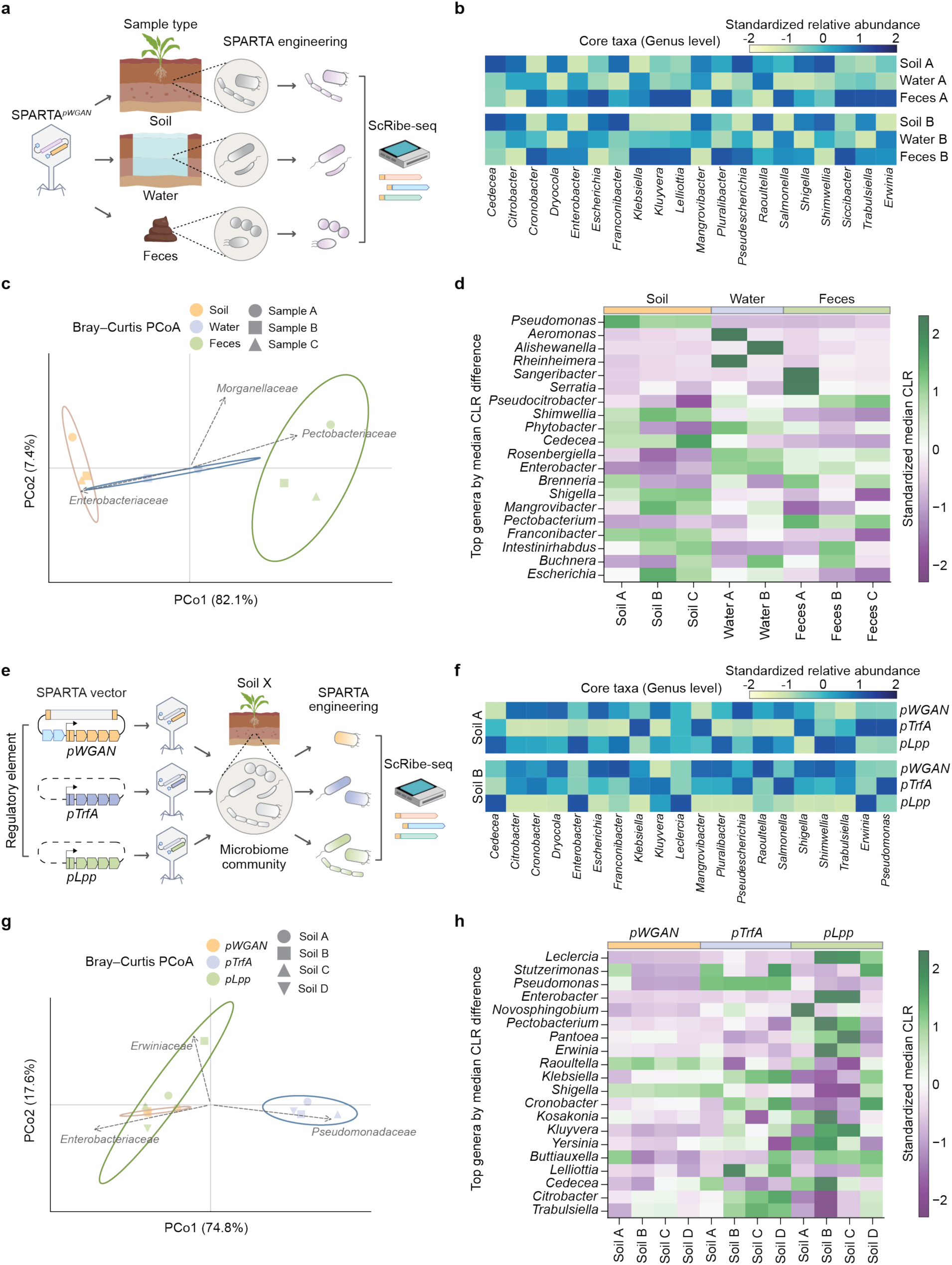
Versatility and modularity of the SPARTA system for *in situ* engineering of native microbiome consortia. **(a)** Overview of SPARTA engineering applied to diverse environmental sample types. Soil and water samples were collected from natural field sites and fecal samples were obtained from specific pathogen free (SPF) mice. **(b)** Standardized relative abundance of representative ScRibe-seq taxa (genus level) across different types of samples engineered by SPARTA. Representative taxa were defined within each sample type as those with the highest mean relative abundance across all conditions. **(c)** Principal coordinate analysis (PCoA) of ScRibe-seq taxa based on Bray-Curtis distance comparing SPARTA engineered microbiome across sample types. **(d)** Heatmap of median centered log ratio (CLR) difference of ScRibe-seq taxa at the genus level across 8 samples with three different sample types (top 20 taxa by highest median CLR difference). **(e)** Design of SPARTA particle variants containing different regulatory elements of the CAST machinery aiming to modulate the engineering profile in soil samples. All particle variants were programmed to target UTS3 and applied in three sequential doses (T3) to soil samples. Microbiome engineering outcomes were analyzed with ScRibe-seq after 3 days of static incubation. **(f)** Standardized relative abundance of representative ScRibe-seq taxa (genus level) across SPARTA conditions for each soil sample. Representative taxa were defined as those with the highest mean relative abundance across all conditions within a given soil sample. **(g)** Principal coordinate analysis of ScRibe-seq genus profile based on Bray-Curtis distance resulting from soil microbiome engineering with SPARTA variants containing different regulatory elements for each soil sample. **(h)** Heatmap of median CLR difference of ScRibe-seq taxa at the genus level across the four soil samples and three regulatory element designs (top 20 genera by highest median CLR difference). The ellipse borders represent 95% confidence interval and dashed arrows are representative correlation vectors for taxa with higher prevalence (>0.5) assessed by envfit analysis.

Additional independent analysis was performed to further validate engineered taxa detected in low prevalence or relative abundance. ScRibe-seq reads from all SPARTA treated samples were analyzed with Emu^41^, a nanopore-tailored classifier providing species-level resolution. This analysis reported 59 engineered species spanning 16 families, including previously identified sample-specific taxa such as *Aeromonas*, *Alishewanella*, and *Rheinheimera* (**Extended Data Fig. 4**). We also detected engineering lineages beyond the canonical *Gammaproteobacteria E. coli* P1 host boundaries, evidencing cross-class, cross-phylum events and possibly beyond (**Extended Data Fig. 4**). This result suggests that SPARTA can mediate chromosomal integration across considerable phylogenetic distances albeit at relatively low engineering abundances. Altogether, SPARTA showcases the ability to engineer diverse microbial consortia across a wide variety of different environmental samples.

### Genetic modularity of SPARTA controls patterns of engineering profiles

SPARTA engineering followed by ScRibe-seq allows us to monitor and interpret engineering outcomes as a function of genetic designs. Engineered relative abundance reported from ScRibe-seq provides guidelines to compare design variants of SPARTA by assessing how individual components govern the overall dynamics of consortia engineering. To demonstrate the genetic modularity of the SPARTA system, we constructed different SPARTA vectors by modifying the promoters for CAST proteins and crRNA expression (**Fig. 3e)**. Because the original promoter (*pWGAN*) was previously optimized for *E. coli*^42^, we evaluated alternative promoters, *pTrfA* and *pLpp*, which are expected to have a broader host expression. *pTrfA* drives the *trfA* gene that initiates replication at the *oriV* of the broad–host-range plasmid *RK2*^43^, suggesting activity across diverse hosts. *pLpp* controls transcription of lipoprotein gene *Lpp*, which has been characterized as having ubiquitous and effective expression among various species in the *Enterobacterales* order^34^.

SPARTA engineering of soil microbiome with promoter variants modulated relative abundance patterns of *in situ* genomic engineering. ScRibe-seq yielded promoter-specific engineering profiles with shifts in the relative abundance of engineered genera despite identical environmental contexts (**Fig. 3f** and **Extended Data Fig. 5a**). SPARTA engineered taxa were clustered primarily by promoter variant rather than by soil of origin (**Fig. 3g**; PERMANOVA, *P* = 0.0026, *R²* = 0.7586), suggesting the regulatory elements comprise a key module reshaping the population targeted by SPARTA. Vector fitting analysis (envfit) further demonstrated that SPARTA vectors with *pTrfA* achieve higher engineered abundance within the *Pseudomonadaceae* family than other regulatory variants. This trend was also observed by CLR comparison, highlighting the enrichment of *Stutzerimonas* and *Pseudomonas* (both under the *Pseudomonadaceae* family) through *pTrfA* SPARTA vectors, as compared to the other variants (**Fig. 3h**). Additionally, the *pLpp* SPARTA variant yielded the highest alpha diversity of SPARTA integrants (**Extended Data Fig. 5b)**, likely attributable to higher engineered abundance across multiple *Enterobacterales* taxa which is expected to have higher delivery efficiency with SPARTA particles. These results agree with differential analysis directly comparing the relative abundance of taxa across promoter variants (**Extended Data Fig. 5c**).

This taxonomic analysis of ScRibe-seq data resulting from SPARTA vector variants demonstrates that reprogramming regulatory elements can redirect SPARTA engineering towards specific lineages (*e.g.*, *Pseudomonadaceae* with *pTrfA*) or broaden engineering coverage (*e.g*., *Enterobacterales* with *pLpp*). These findings highlight SPARTA as a modular, programmable platform that can be rationally tuned at the genetic design level, translating choices in promoter architecture into significant shifts in community-level engineering outcomes. By directly coupling genotype-to-phenotype mapping through ScRibe-seq, SPARTA advances toward a predictable and customizable chassis for precision *in situ* microbiome engineering of native samples to achieve desired community compositions and functions.

### *In situ* delivery and expression of DNA cargo through SPARTA

SPARTA enables isolation of individual genetically reprogrammed native bacterial strains from soil samples. While ScRibe-seq analysis delineates the spectrum of bacterial cell types potentially engineered via SPARTA, it alone is not sufficient to prove that the recipient cells are capable of expressing integrated DNA cargos. To validate this in individual members of native communities, we constructed a tandem reporter cassette encoding a bioluminescence reporter and an antibiotic resistance marker (**Fig. 4a**). Integration of the cargo enables both phenotypic detection and selective enrichment post-engineering. After SPARTA treatment of soil, we isolated multiple resident bacterial strains expressing the engineered cargo DNA, confirmed through bioluminescence phenotype screening (**Fig. 4b**) and whole genome sequencing (WGS) (**Fig. 4c**). The recovered isolates spanned multiple genera across *Enterobacteriaceae*, *Erwiniaceae*, and *Pseudomonadaceae*, consistent with the ScRibe-seq–derived engineering profile. The isolated representative species exhibited different levels of bioluminescence intensities in phenotyping assays, suggesting species-dependent differences in transgene expression (**Fig. 4b**). All assembled genomes exhibited cassette insertions at one of the designated 16S rRNA target sites (**Fig. 4c** and **Extended Data Fig. 6a**), confirming SPARTA-dependent genome integration events.

**Figure 4.**
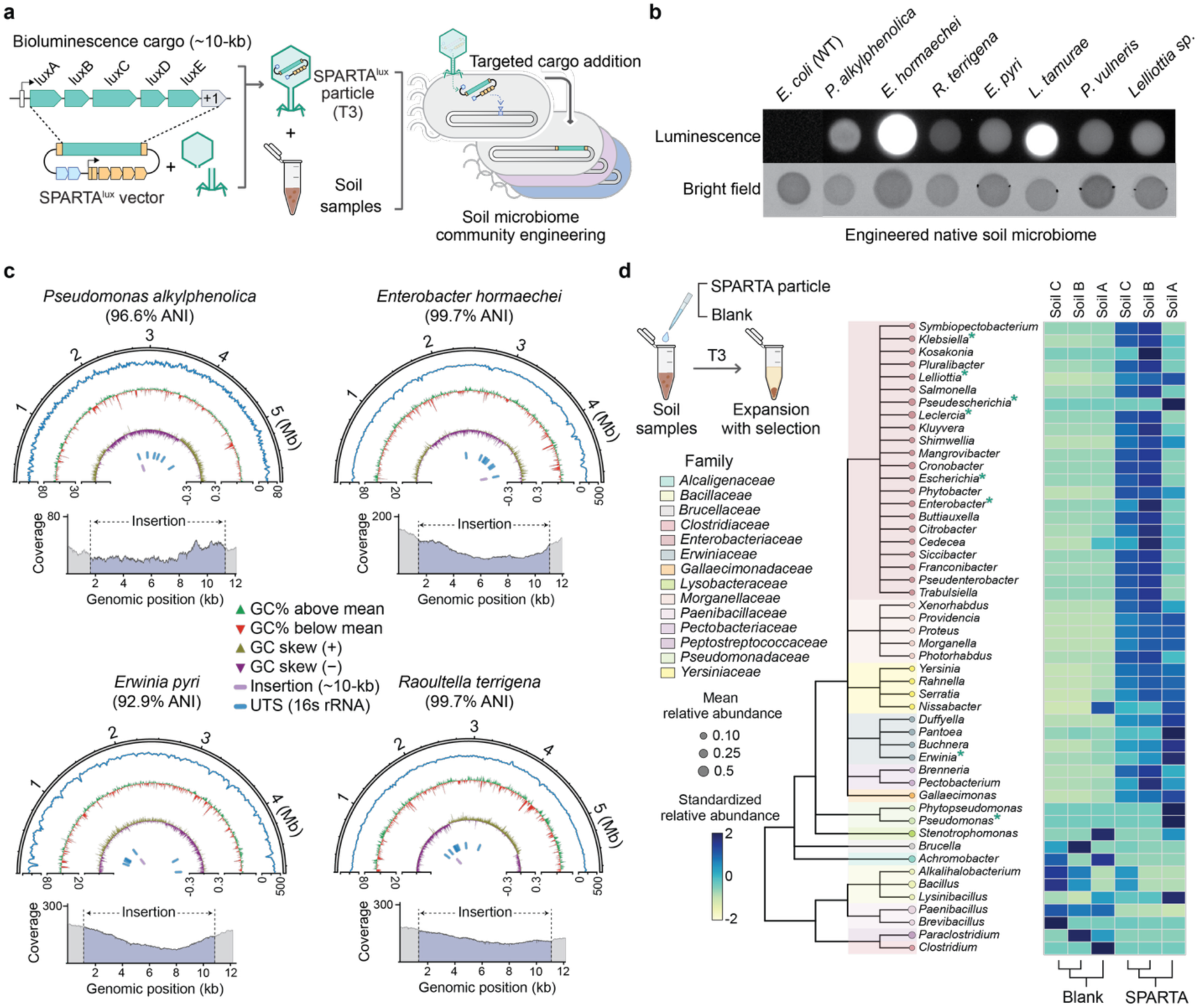
Validation of functional cargo engineering and expression in SPARTA-treated native soil microbiomes. **(a)** Bioluminescence genes carrying a positive selection marker (+1) were encoded as cargos within the SPARTA^lux^ vector and delivered into soil samples with three treatments (T3). **(b)** Representative luminescent phenotype spotting assay on selective agar plates showing bioluminescence expression in wild-type *E. coli* (WT) control and multiple native bacterial genera engineered by SPARTA^lux^ particles. **(c)** Validation of SPARTA-directed genome editing in isolated soil bacterial species using whole-genome sequencing. Semicircular plots display read coverage, GC content (%), and GC skew (from outer to inner rings) along the genomic position and detailed coverage plot of the cargo insertion site. Blue and purple features indicate the target loci (16S rRNA) and the cargo integration site, respectively. ANI denotes average nucleotide identity estimated by sourmash. **(d)** To compare community enrichment, soil samples were treated with blank solution or SPARTA particles (T3) prior to expansion. Standardized relative abundance of bacterial genera in cultures of soil microbiomes after enrichment with antibiotics. Clade and node colors represent family-level taxonomy, and node size indicates the mean relative abundance of taxa across samples.

SPARTA allows community-level enrichment of engineered populations in agreement with ScRibe-seq readout. Motivated by robust cargo expression across phylogenetically diverse native species, we enriched the engineered bacteria community by co-culturing the SPARTA-treated soil under antibiotics selection. Although soil cultures are subject to strong biases due to growth conditions and intrinsic antibiotic resistance in microbes, 16S rRNA gene amplicon sequencing revealed preferential enrichment of taxa from *Gammaproteobacteria* class, including *Enterobacteriaceae*, *Erwiniaceae*, *Morganellaceae*, *Pseudomonadaceae*, and *Yersiniaceae,* in SPARTA treated soil as compared to untreated controls (**Fig. 4d** and **Extended Data Fig. 6b**). The enriched populations overlapped extensively with those detected by ScRibe-seq (**Extended Data Fig. 6c**), indicating that engineered members can be selectively propagated post-engineering. Taken together, these results validate the precise genetic modulation and expression of synthetic DNA within native microbiomes via SPARTA. Furthermore, the strong correlation between ScRibe-seq result and selectively enriched taxa demonstrates that ScRibe-seq accurately represents the engineering profile of bacterial consortia.

### SPARTA enhances fecal microbiome transplantation

We next systematically explored the possibility of SPARTA-enhanced fecal microbial transplantation (FMT) as a clinical application. FMT has emerged as a powerful therapeutic strategy for a wide range of microbiome-associated diseases^44,45^, including inflammatory bowel disease^46^, metabolic syndromes^47^, neuropsychiatric disorders^48^, and responses to cancer immunotherapy^49^. Recent discoveries underscoring the diverse roles of the gut microbiome on human physiology suggest that FMT could be extended far beyond gastrointestinal health^50^. Unlike conventional *ex vivo* manipulations, SPARTA introduces defined genetic functions into a target chassis *in situ* without perturbing the native community. This strategy preserves donor-derived microbial diversity while enabling integration of engineered traits. Consequently, donor microbiome – containing taxa that have stably acquired exogenous functional DNA in their genome via SPARTA – can be transferred intact into recipient hosts (**Extended Data Fig. 7a**). By designing genetic cargos to improve therapeutic efficacy, SPARTA-coupled FMT offers opportunities for next-generation, programmable, and personalized microbiome therapeutics.

SPARTA engineering during the FMT process was successfully characterized by using a murine fecal model. Although multiplexed engineering in murine fecal samples was demonstrated above (**Fig. 3**), *E. coli* was chosen as a safe and tractable chassis for *in vivo* enrichment and heterologous expression of exogenous DNA. After donor mice were colonized with *E. coli* strains, fecal samples were treated with SPARTA^lux^ particles targeting the *prfC* gene (T2 dosage) and were subsequently transplanted into recipient mice (**Fig. 5a**). To evaluate parameters governing initial engraftment and long-term persistence, we compared several antibiotic treatment regimens, including *ex vivo* selection (applied to donor feces during engineering) and *in vivo* selection (administered to recipients post-FMT). Successful transfer and engraftment of SPARTA engineered cells was confirmed by luminescence imaging of recipient stool samples one day post-transplantation (**Fig. 5b** and **Extended Data Fig.7b**). The population of engineered cells was subsequently monitored by measuring CFU of integrants in recipient feces over time. While both *ex vivo* and *in vivo* selection schemes yielded comparable initial engraftment, *in vivo* selection exhibited enhanced long-term FMT-derived CFU persistence up to 35 days post-FMT (**Fig. 5c**). Additionally, transposon-end sequencing analysis revealed high insertion accuracy (>95%) among integrants stably maintained for over a month (**Fig. 5d**). Together, these results demonstrate that SPARTA *in situ* engineering can be coupled with FMT to achieve stable transfer and persistence of engineered donor microbiome in recipients.

**Figure 5.**
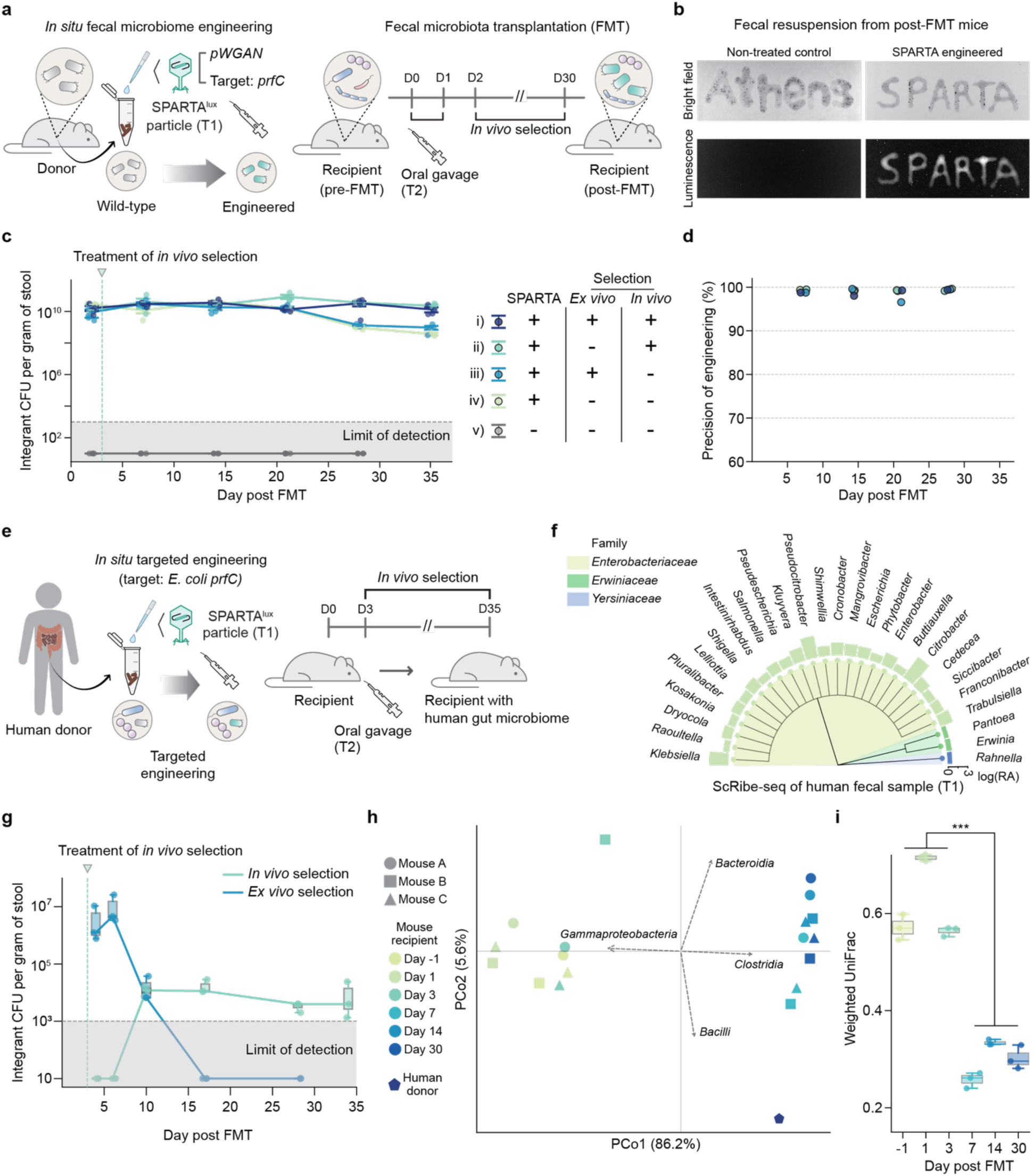
Fecal microbiome transplantation enhanced by SPARTA engineering. **(a)** Schematic of *in situ* fecal microbiome engineering by targeting *E. coli* cells colonized in a donor mouse model, followed by fecal microbiome transplantation to recipient mice. SPARTA^lux^ particles designed with the *pWGAN* promoter and the *prfC* targeting crRNA were used to engineer *E. coli* in donor feces (T1 treatment). Engineered fecal suspensions were administered to recipients by oral gavage once daily for two consecutive days, and recipient microbiomes were monitored for one month post-FMT. **(b)** Representative bright field and luminescence images of resuspended stool samples collected from mice transplanted with non-treated or SPARTA engineered *E. coli* cells. **(c)** Population of engineered *E. coli* cells in FMT recipients over 5 weeks post-FMT under different antibiotics selection schemes. *In vivo* selection indicates antibiotic treatment via drinking water and *ex vivo* selection indicates antibiotics included in the phage particles resuspension during SPARTA engineering. Error bars indicate the mean ± standard error of the mean (SEM) from three experimental replicates (N = 3). The vertical dashed line indicates the start of antibiotic treatment for *in vivo* selection, while the horizontal dashed line and gray shaded area below indicate the limit of detection. **(d)** Precision of SPARTA engineering assessed by transposon-end sequencing using DNA extracted from recipient fecal samples. **(e)** Schematic of FMT using SPARTA-engineered human stool to transplant SPARTA engineered human gut microbiome in a mouse model. SPARTA^lux^ particles (*pWGAN* promoter and *prfC* target) were employed for targeted *E. coli* engineering in human fecal sample (T1). Engineered human stool resuspensions were administered to recipients by oral gavage on two consecutive days, and the recipient microbiomes were tracked for one month post-FMT. **(f)** Genus-level taxonomic profile (ScRibe-seq) of SPARTA (UTS, T1)-treated human stool, displayed on a phylogenetic tree colored by family. Bars indicate the relative abundance of each taxon. **(g)** Population of engineered cells (luminescence-positive) in post-FMT recipient stool quantified as CFU/g of feces. The vertical dashed line indicates the start of antibiotic treatment for *in vivo* selection, while the horizontal dashed line and gray shaded area below indicate the limit of detection. **(h)** PCoA analysis based on weighted UniFrac comparing 16S rRNA profiles from recipient (*in vivo* selection) and human donor stools. Dashed arrows show representative correlation vectors for taxa (class level) with prevalence >0.5 assessed by envfit analysis. **(i)** Weighted UniFrac dissimilarity of microbiome composition between recipient mice and the human donor. Three asterisks indicate P <0.001, assessed by one-way ANOVA. All boxes span from the 25th to 75th percentiles (interquartile range) from three experimental replicates (N = 3), whiskers extend to 1.5 times the interquartile range and center lines represent the median.

### *In situ* engineering of human microbiome and transplantation into a murine model

SPARTA-enhanced FMT can be extended to clinically relevant and compositionally complex human microbiomes. To test this, we next performed SPARTA-mediated FMT with human donor fecal samples (**Fig. 5e**). Targetable taxa within human feces were first identified using ScRibe-seq following treatment with universal targeting SPARTA particles (UTS, T1) (**Fig. 5f**). The resulting taxonomic profile revealed the presence of *Escherichia-Shigella* clade, which is considered as a safe and suitable target for SPARTA engineering. Based on this analysis, native *E. coli* cells within the human donor gut microbiome were targeted using SPARTA particles programmed to integrate at the *prfC* locus (**Fig. 5e**). Analogous to the murine model FMT experiments presented above, we applied both *ex vivo* and *in vivo* selection conditions to compare the dynamics of engineered populations following transplantation into mice. As expected, the lower baseline relative abundance of *E. coli* and greater compositional variability in human fecal material reduced both initial engineering efficiency and early engraftment (within 3–5 days post-FMT; **Fig. 5g**). Despite this, under *in vivo* selection, luminescent *E. coli* colonies emerged after day 7 and persisted at stable levels for up to 30 days post-FMT, whereas *ex vivo* selection populations declined and became undetectable by ∼20 days. Engineered human native *E. coli* colonies were isolated, and sequence-verified by WGS, validating precise engineering in a desired location of multiple native *E. coli* strains (Extended Data Figs. 8a–b).

A SPARTA engineered human fecal microbiome was successfully engrafted into the mouse gut. To assess the broader community-level composition of human-to-mouse microbiome transfer, we profiled the fecal microbiomes of recipient mice over time. Principal coordinate analysis (PCoA) revealed a significant compositional shift of the recipient microbiome community toward that of the human donor, particularly pronounced after day 7 post-FMT (**Fig. 5h**; PERMANOVA, *P* = 0.0001, *R²* = 0.9652 and **Extended Data Fig. 8c**). This trend was also confirmed by UniFrac distance analysis^51^ (**Fig. 5i)**, demonstrating sustained similarity between recipient and donor communities through day 30. Collectively taken, these findings establish that SPARTA enables stable genomic integration and transfer of human-derived taxa during FMT, leading to the formation of a murine gut ecosystem harboring synthetically modified human bacteria populations. It highlights the feasibility of generating synthetically engineered microbiomes through SPARTA in fecal samples for downstream transplantation.

## Discussion

Our work introduces SPARTA as a novel system for *in situ* native microbiome reprogramming in diverse environments with minimal disturbance of taxonomic composition. This strategy of SPARTA allowed us to synergistically pioneer ScRibe-seq as the first high-throughput readout system for direct taxonomic classification and quantification of engineered microbial consortia in native samples.

Precise cargo integration into the 16S rRNA genome using SPARTA enables ScRibe-seq to selectively amplify the modified genome from complex, noisy metagenomic samples. By directly reading out 16S rRNA sequences of only engineered taxa, ScRibe-seq captures unbiased engineering profiles in a single step, which is challenging to achieve with conventional sequencing approaches that rely on multi-stage enrichment of engineering signal. This targeted readout strategy further obviates the need for additional whole-metagenome sequencing, thereby improving the scalability and resolution of the analysis. Furthermore, high sensitivity of ScRibe-seq facilitates the identification of previously unrecognized host–phage interactions beyond what has been reported in existing literature. We hypothesized this is due to P1’s native host range inherently being constrained by a need for phage replication that does not apply to SPARTA particles. In this work, we circumvent these limitations and unlock our ability to both engineer native microbiomes and to detect, classify, and quantify engineering events.

The engineering dynamics and host spectrum of SPARTA were shown to be controllable by modularizing SPARTA vector components. The programmability of SPARTA is derived from multiple genetic components, including regulatory elements of the CAST system, its cognate crRNA target, and structural genes (e.g., tail proteins) of phage particles. The choice of selection markers within a cargo DNA also affects the enrichment profiles of engineered populations and enables the isolation of specific individual species. Achieving both specificity and generality in a microbiome genome engineering system is inherently challenging and seemingly paradoxical. By encoding multiple different layers of programmability, our findings suggest the possibility of flexible community engineering through SPARTA, either broadening targeting or restricting it to defined microbiome clades. Future integrating this design space with artificial intelligence-guided modeling^52^ and genome language model-based prediction^53^, the engineering range of SPARTA can be further expanded toward previously inaccessible bacterial clades that play critical ecological or functional roles in their habitat. Subsequent designs and findings could greatly increase the breadth, precision, and efficiency of SPARTA as a versatile *de novo* programmable native microbiome editing platform in environmental samples.

SPARTA demonstrated successful reprogramming of resident microbiomes regardless of physical or chemical properties of the habitat, highlighting its compatibility with diverse environmental contexts. Unlike traditional approaches that rely on introducing exogenous or laboratory-adapted bacterial strains, which often suffer from poor ecological fitness, our method enables direct genetic modification of resident microbial hosts that are already evolutionarily adapted to their ecological niches. Additionally, targeted chromosomal integration of DNA payloads through SPARTA ensures stable expression of synthetic functions. Overall, we demonstrate that SPARTA circumvents the long-standing bottleneck of establishing foreign strains in complex ecosystems and opens a path toward long-term and sustainable engineering of wild microbiome in their native environment.

In short, we designed, applied, and readout SPARTA to demonstrate its potential as a generalizable platform for manipulating communities of environment-resident microbiomes *in situ* by installing synthetically designed and constructed genetic information. SPARTA establishes a foundation to enable robust, long-term native microbiome remodeling across environmental, agricultural, and clinical applications.

## Methods

### Strains and plasmids construction

All bacterial strains used in this study are listed on **Supplementary Table 1**. *Agrobacterium tumefaciens* was purchased from GoldBio. *Shigella flexneri* (CFS100) was kindly provided by Shelley M Payne, University of Texas at Austin, as a gift. All other bacterial strains were acquired from ATCC. For *R6K* plasmid construction and DNA integration screening, *E. coli* DH10B (Δ*YahA::piR-cmR*) strain was generated through recombination by using the previously constructed recombination plasmid, pKW20. The phage packaging strain C600 containing *P1kc* Δ*coi* (a gift from Nathan Crook, North Carolina State University) was further engineered to generate synthetic P1 genomes, such as *P1kc Δcoi:piR* and *P1kc Δcoi:piR pacA*\*, through recombination. These modified lysogens were used to manufacture phage particles with *R6K* based construct and to increase the yield of recombinant phage particles.

All SPARTA vectors were constructed using Gibson or Golden gate assembly methods. Spacer gRNA fragments were generated by annealing two complementary DNA oligonucleotides and subsequently ligating them into SPARTA backbones via Golden Gate assembly. DNA oligonucleotides were ordered through Integrated DNA Technologies (IDT) or MilliporeSigma, while PCR fragments for construction were amplified using PrimeSTAR GXL (Takara) or repliQa HiFi polymerase (Quantabio). CAST and phagemid components were derived from pSL1777 (Addgene #160731) and BBa_J72214-BBa_J72090 (Addgene #40782), respectively. Additional DNA fragments for building SPARTA vectors were custom-synthesized through Twist Bioscience and incorporated using Gibson assembly. All constructs were made in *E. coli* DH10B and DH10B (Δ*YahA::piR-cmR*) cells and listed in **Supplementary Table 1**.

### Growth conditions of laboratory strains

Bacteria cells were cultured in Luria-Bertani (LB) medium or on LB-Agar plates containing appropriate antibiotics at either 30°C or 37°C. For selection and genetic engineering of *Enterobacteriales* bacteria and *A. tumefaciens*, kanamycin (50 µg/ml) and gentamycin (20 µg/ml) were used; for *P. putida*, tetracycline (10 µg/ml) was used. The SPARTA phage packaging strain was cultivated in the presence of kanamycin or chloramphenicol.

### Phenotyping assays and genotyping PCR analysis

Individual colonies grown on selective LB-agar plates were picked and resuspended in 60 μl of double-distilled water (ddH_2_O). For phenotyping, 4 μl of cell suspension was spotted onto selective agar plates for 18 hours at either 30°C or 37°C incubation, depending on the strain and assay. For genotyping, aliquots of the colony suspensions were used as PCR templates to confirm targeted DNA insertions within the target locus of the bacterial genome. PCR reaction was carried out using PrimeSTAR GXL (Takara) or repliQa HiFi (Quantabio) polymerase. For gel analysis, 2 µl of each PCR reaction was mixed with 5.5 µl ddH₂O and 1.9 µl 6× gel loading dye (New England Biolabs) and resolved on 1% (w/v) agarose gels containing SYBR Safe DNA stain (Thermo Fisher Scientific). Electrophoresis was performed at 100 V for 30 min, and DNA bands were visualized using a ChemiDoc Imaging System (Bio-Rad).

### Phage particle generation and titer measurement

The P1 lysogenic strain, *E. coli* C600-P1kc (Δ*coi KanR::piR-cmR)*, was transformed with the SPARTA vectors via electroporation or bacterial conjugation. Phage particle production and transduction was performed according to previously established protocols with minor modifications. Briefly, overnight cultures of the packaging strain were sub-inoculated (1:100 dilution) into phage generation medium consisting of LB media supplemented with 100 mM MgCl_2_ and 5mM CaCl_2_, together with the appropriate antibiotics. The strain was induced when the culture reached an OD_600_ value of 1.0. After an additional 2 hours of incubation, phage lysates were treated with chloroform (1:40 v/v) for 15 minutes on ice and then centrifuged at 3,000–4,000 g-force for 10 minutes. The supernatant containing phage particles was collected into new tubes and stored at 4°C until use. If needed, phage particles were buffer-exchanged into phage resuspension buffer (50 mM Tris−HCl, 8 mM MgSO4, and 100 mM NaCl, 100 mM MgCl2, 10 mM CaCl2, pH 7.5), which was also used for phage dilutions to adjust desired titers. All phage stocks were used within six months of preparation.

For titer evaluation of isolated SPARTA particles, recipient *E. coli* cells (DH10B or DH10B Δ*YahA::piR-cmR*) were grown overnight and sub-inoculated into phage generation medium at a 1:100 dilution. When cultures reached OD_600_ of 1.0, cell suspensions were mixed with phage particles at a 3:1 volume ratio and co-incubated at 30°C or 37°C for 20 min with shaking, followed by 20 min under static conditions. Transduction was quenched by adding SOC media supplemented with 200 mM sodium citrate (1/4 volume of the total reaction) and incubating for an additional 40 minutes at 30°C or 37°C. Transduction efficiency (phage titer) was determined by plating on LB agar containing the appropriate antibiotics.

### *In vitro* engineering via SPARTA

Recipient cells were cultured overnight in LB medium at 30°C or 37°C and then sub-inoculated into phage generation medium for SPARTA particle transduction as described above. Following transduction, cells were incubated at room temperature or 30°C for an additional 1–2 hours to facilitate DNA integration mediated by the CAST system. Cultures were then sub-inoculated into fresh LB medium containing kanamycin or tetracycline and incubated overnight to recover and enrich integrant populations. Integration efficiency (CFU) and genotype verification of colony isolates were assessed by plating on selective LB agar according to the cargo. PCR genotyping of individual colonies was performed as needed to confirm successful integration.

SPARTA vectors based on R6K origin were used for *E. coli* and CDF backbone constructs were employed for *P. putida* and *A. tumefaciens*, was employed to enable iterative cargo integration. The *A. tumefaciens* SPARTA construct incorporated a *pTet* inducible promoter to enhance expression of CAST effector proteins, whereas constructs for other species used the constitutive *pWGAN* promoter. Iterative engineering was performed following the same SPARTA engineering protocol, with supplementation of 200 ng/ml anhydrotetracycline (ATC) to induce CAST expression in *A. tumefaciens* during SPARTA particle transduction. Iterative engineering following the engineering protocol described beforehand, with 200 ng/ml anhydrotetracycline (ATC) inducer supplementation into the *A. tumefaciens* culture during phage particle transduction. This process was repeated for two additional rounds to obtain the final engineered strains. All resultant strains were sequence-verified by amplicon sequencing or whole-genome sequencing (Nanopore). Genomic target sites for all three integration rounds and primer sequences used for PCR genotyping are detailed in **Supplementary Tables 2** and **5**.

### Environmental samples collection

Soil and water samples were collected on the California Institute of Technology (Caltech) campus courtyard at the indicated GPS coordinates, as detailed in **Supplementary Table 3**. After collection, samples were weighted and transferred into microcentrifuge tubes. All samples were processed for the experiment on the day of collection to preserve the original microbiome composition.

### *In situ* engineering via SPARTA

For soil samples, intact material was used directly for SPARTA engineering without additional homogenization step to simulate the field application. Samples were transferred to microcentrifuge for weighing and then incubated statically with 200 µl of blank resuspension buffer (50 mM Tris−HCl, 8 mM MgSO4, and 100 mM NaCl, 100 mM MgCl2, 10 mM CaCl2, pH 7.5) or with the same buffer containing either naked SPARTA vector DNA or SPARTA particles. Incubation proceeded for 24 hours at room temperature for each treatment. After incubation, solid samples were washed twice with a PBS buffer (liquid samples were not washed) and collected for downstream analyses, such as DNA extraction, enrichment culture, plating, or bacterial isolation*. In situ* engineering of water samples was processed as described above without the PBS washing step.

### *In situ* sample processing and DNA extraction

All environmental samples were cultured on the brain heart infusion (BHI) broth or on BHI-agar plate at 30°C or 37°C, in the presence of antibiotics (kanamycin 50 µg/ml and tetracycline 10 µg/ml) supplemented with cycloheximide to inhibit fungal growth. DNA from enriched microbial communities or isolated colonies was extracted using the DNeasy Blood & Tissue Kit (Qiagen). For direct DNA extraction from *in situ* samples, such as soil, water, or fecal material, the DNeasy PowerSoil, PowerWater, and QIAamp PowerFecal kits (Qiagen) were used, respectively.

### NGS analysis

Synthetic DNA insertion via SPARTA was validated through amplicon sequencing or whole genome sequencing by using nanopore sequencing on R 10.4.1 flow cells (Oxford Nanopore Technologies). Amplicons were generated with primers annealing to the sequences flanking targeted integration sites and then prepared by Ligation Sequencing Kit V14 SQK-LSK114 (Oxford Nanopore Technologies). For validation of native strains engineered by SPARTA, genomic DNA (gDNA) extracted with the DNeasy Blood & Tissue Kit (Qiagen) was used for nanopore whole genome sequencing.

To prepare transposon sequencing library, gDNA was extracted with DNeasy Blood and Tissue kits (Qiagen) using the bulk cell populations after cargo DNA integration via SPARTA. Libraries were generated with the Illumina DNA Prep tagmentation kit following the manufacturer’s protocol, using custom primers targeting the transposon arm (R-end) together with standard Illumina sequencing primers compatible with the kit. Sequencing was performed on an Illumina MiSeq instrument using custom read primers. Library concentrations were quantified with the Qubit 1× dsDNA High Sensitivity Assay Kit (Invitrogen, Thermo Fisher Scientific) for normalization prior to sequencing. All primers used for amplicon sequencing and transposon sequencing are listed in **Supplementary Table 5**.

### ScRibe-seq analysis

For ScRibe-seq library preparation, DNA extracted directly from *in situ* samples was amplified with PCR reaction using Q5® High-Fidelity (New England Biolabs) with primers designed to generate long amplicon (∼1,100–1,400-bp) spanning the transposon end and near full length 16S rRNA, including custom barcode sequences. Amplifies products were purified and size-selected by gel extraction using the QIAquick PCR purification and gel extraction kits (Qiagen). Then libraries were prepared with the Ligation Sequencing Kit V14 (SQK-LSK114) (Oxford Nanopore Technologies) and normalized by concentration as measured with the Qubit 1× dsDNA High Sensitivity Assay Kit. Sequencing was conducted on R 10.4.1 flow cells using a MinION Mk1C (Oxford Nanopore Technologies). During sequencing, reads shorter than 800-bp were excluded to achieve high-depth coverage (∼1 million reads per sample) of target long amplicons. Raw reads were processed by quality filtration, denoising, adapter trimming, and chimera detection to obtain the high quality ScRibe-seq reads for downstream metagenomic analysis.

### 16S rRNA metagenomic analysis

Conventional full-length 16S rRNA gene amplicon sequencing was also performed with R 10.4.1 flow cells (Oxford Nanopore Technologies) and then denoised by Nanoclust^54^. Taxonomic classification of metagenomic reads, including conventional 16S rRNA and ScRibe-seq data, was done with Kraken2 and Bracken based on NCBI 16S RefSeq database. Taxa with relative abundance below 0.05% at the genus level were excluded for downstream analysis unless otherwise noted. For detection of low abundant taxa, Emu taxonomic classification was employed with minimum relative abundance threshold of 0.03% or a total read count of 50. Taxonomic lineage assignments were based on the NCBI Taxonomy and used for phylogenetic tree construction. Differential abundance analysis of taxonomic profiles was performed with ALDEx2, using Monte Carlo sampling from the Dirichlet distribution as described previously. Dimensionality reduction analyses were based on Bray–Curtis and weighted UniFrac distances to assess beta-diversity among metagenomic datasets. The phylogenetic tree for weighted UniFrac analysis was constructed from NCBI Taxonomy with MUSCLE alignment.

### Mice

Wild-type specific pathogen free (SPF) C57BL/6J (Jackson Laboratories, RRID:IMSR_JAX:000664) males and females were used for experiments. Mice were 8 weeks old at the start of all experiments. Upon receipt, mice were acclimated to the facility for at least 5 days prior to the start of any experiments. Experimental animals were group-housed in sterilized microisolator cages. Mice received autoclaved 5010 PicoLab Rodent Diet (LabDiet) and sterilized water in their home cages *ad libitum*. Ambient temperature in the animal housing facilities was maintained at 21–24°C, 30–70% humidity, with a cycle of 13 hours light, 11 hours dark. All mouse experiments were performed in accordance with the NIH Guide for the Care and Use of Laboratory Animals using protocols approved by the Institutional Animal Care and Use Committee at the California Institute of Technology.

### *In situ* murine fecal microbiome engineering and transplant

To colonize the donor mice *E. coli* MG1655 were resuspended in the gavage buffer (1.5% NaHCO_3_ in PBS). 200 µl of resuspension, containing 10^8^–10^9^ CFUs, was administered to each mouse through oral gavage. To assist with colonization, streptomycin sulphate (5 mg/ml, GoldBio) was added to the drinking water of the donor mice seven days prior to the E. coli oral gavage. Once a stable *E. col*i colonization was confirmed, stool samples were collected from the donor mice, combined, and 50 mg of the mix was co-incubated with SPARTA particle resuspensions for 24 hours (T1). The engineered fecal sample was washed twice with PBS solution and then resuspended in 200 µl of gavage buffer. Some of the samples were treated with kanamycin monosulphate (50 µg/ml) and tetracycline hydrochloride (10 µg/ml, Millipore Sigma) for *ex vivo* selection of engineered microorganisms. The samples were washed with PBS and resuspended in the gavage buffer. They were administered to the recipient mice via single oral gavage. To deplete the host microbiome and facilitate colonization with the engineered taxa, where specified, the recipient mice were treated with the mixture of antibiotics in drinking water: vancomycin hydrochloride (0.5 mg/ml, Medisca), erythromycin (0.01 mg/ml, Fisher Scientific), neomycin (1 mg/ml, Fisher Scientific), gentamicin sulphate (0.1 mg/ml, GoldBio), and ampicillin sodium (1 mg/ml, GoldBio) (VENGA mix). The antibiotic treatment was performed for 10 days before the delivery of the engineered fecal samples. Recipient mice were transferred back to the regular water 2 days before the delivery of the fecal samples. After the engineered fecal sample was delivered into the recipient, mice were provided with drinking water supplemented with streptomycin sulphate (5 mg/ml), kanamycin monosulphate (0.4 mg/ml, GoldBio), and tetracycline hydrochloride (0.03 mg/ml). Stool samples of recipient mice were collected up to 35 days post-FMT to monitor the persistence of SPARTA-engineered E. coli cells. Colonization rates were assessed by fecal sample plating on selective LB agar media.

### *In situ* human microbiome engineering and transplant

Human fecal sample obtained from iSpecimen was resuspended in a deoxygenated phage resuspension buffer, resulting in a 200 mg/ml suspension. Suspension was treated with either blank phage resuspension buffer alone (control samples) or phage resuspension buffer containing SPARTA particles. For *ex vivo* selection, SPARTA-treated samples were then incubated with kanamycin monosulphate (50 µg/ml) and tetracycline hydrochloride (10 µg/ml) for two hours at 37°C to enrich engineered population. Samples were washed with PBS twice, resuspended in the gavage buffer, and 200 µl of a sample was administered to each mouse via three oral gavages over the span of six days (on days D0, D2, and D5). Recipient mice were pretreated with VENGA antibiotic mix for 10 days, then they were transferred to the regular drinking water 24 hours before the first administration of fecal material. On D1 (24 hours after the first gavage), some mice were provided with kanamycin monosulphate (0.4 mg/ml), and tetracycline hydrochloride (0.03 mg/ml) in the drinking water for the *in vivo* selection of the engineered microorganisms. Stool samples of recipient mice were collected up to 35 days post-FMT to monitor the persistence of SPARTA-engineered *E. coli* cells. Colonization rates were assessed by fecal sample plating on selective LB agar media.

### Data and code availability

The sequencing data generated in this study have been deposited in the NCBI BioProject database with the accession number PRJNA1387163. A list of BioProject and SRA accession numbers corresponding to each NGS dataset is described in **Supplementary Table 6**. Scripts used for NGS data analysis are available upon request.

## Acknowledgement

We thank Prof. Nathan Crook from North Carolina State University and Prof. Shelley M. Payne from the University of Texas at Austin for generously providing the *C600-P1kc* strain and *Shigella flexneri CFS100*, respectively. K.W. acknowledges funding support from the Shurl and Kay Curci Foundation, NIH (1DP2GM140937) and Caltech Center for Environmental Microbial Interactions (CEMI). R.J.Z. is supported by NSF (Fellow ID 2020296026). The authors acknowledge Anton Kuzmenko and Elena Fefelova for insightful discussions.

## Author contributions

J.H., C.S. and K.W. conceived this work and designed experiments. J.H., C.S., and B.S. conducted experiments and collected data. B.S. and A.O. carried out animal experiments. J.H., C.S., B.S., M.L., T.G., and R.J.Z generated plasmid constructs and engineered microbial strains used herein. J.H. and Jianyi H. generated NGS libraries. J.H. and C.S. generated computational scripts used for sequencing data analysis and interpreted the results of this work. J.H., C.S., B.S., A.O., K.W., and S.K.M. wrote the manuscript and M.L., T.G., and R.J.Z. participated in discussion and provided feedback.

## Declaration of interests

K.W. is a founder of Genyro Inc.. C.S., R.J.Z., Jinayi H., and K.W. are co-founders of Syntaxa Inc.. Neither company is involved in this study. S.K.M. is co-founder of Axial Therapeutics and Nuanced Health, and has equity in Seed Health, Time BioVentures and Intrinsic Medicine.

---

**Extended Data Figure 1.**
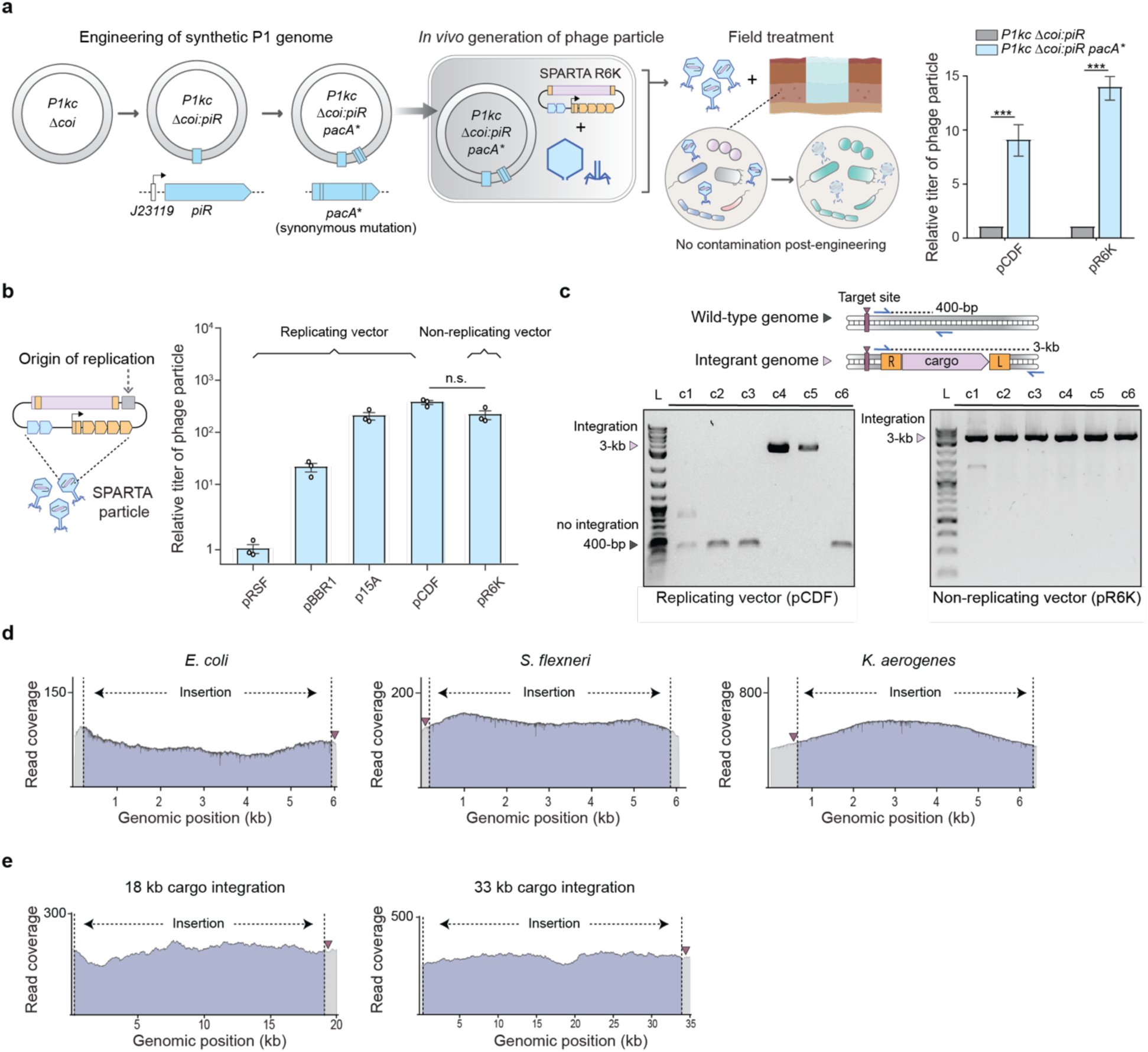
Engineering and optimization of synthetic phage genome and delivery vector for SPARTA. **(a)** Design and engineering of synthetic P1 phage genome for *in vivo* generation of nonvirulent phage-like particles suitable for field applications. The *piR* gene was first inserted into the *P1kc Δcoi* genome to enable replication of *R6K* plasmids, and a synonymous mutation in the *pacA* gene (*pacA*\*) was introduced to replace the wild-type *pacA*. Phage particles were generated from the resultant engineered P1 genomes, minimizing the risk of environmental contamination. Relative titers of particles derived from different synthetic P1 genomes are shown, demonstrating increased yield of particles from the *P1kc Δcoi:piR pacA*\* construct. **(b)** Relative titers of SPARTA particles determined by phage transduction units in *E. coli* cells carrying vectors with different origins of replication. **(c)** Schematic of genotyping analysis of cargo DNA (∼3-kb) integration and representative genotyping results from colony PCR (colonies 1 to 6; c1–6) after engineering with SPARTA vectors based on *CDF* (replicating) and *R6K* (non-replicating) backbones. Blue arrows and dotted lines indicate primer pairs and PCR products, respectively**. (d)** Read coverage of genomic regions including target sites of synthetic DNA integration in different *Enterobacteriales* strains, derived from next-generation sequencing (NGS) data. Representative coverage plots are shown, with purple triangles indicating the integration target sites. **(e)** Read coverage of large cargo integration events (18-kb and 33-kb) in the genome of *E. coli* cells based on NGS data. Purple triangles indicate the target site of integration. All error bars indicate the mean ± standard error of the mean (SEM) from three experimental replicates (N = 3). Three asterisks indicate a P <0.001 and n.s. represents no significant difference, based on the Student’s t-test.

**Extended Data Figure 2.**
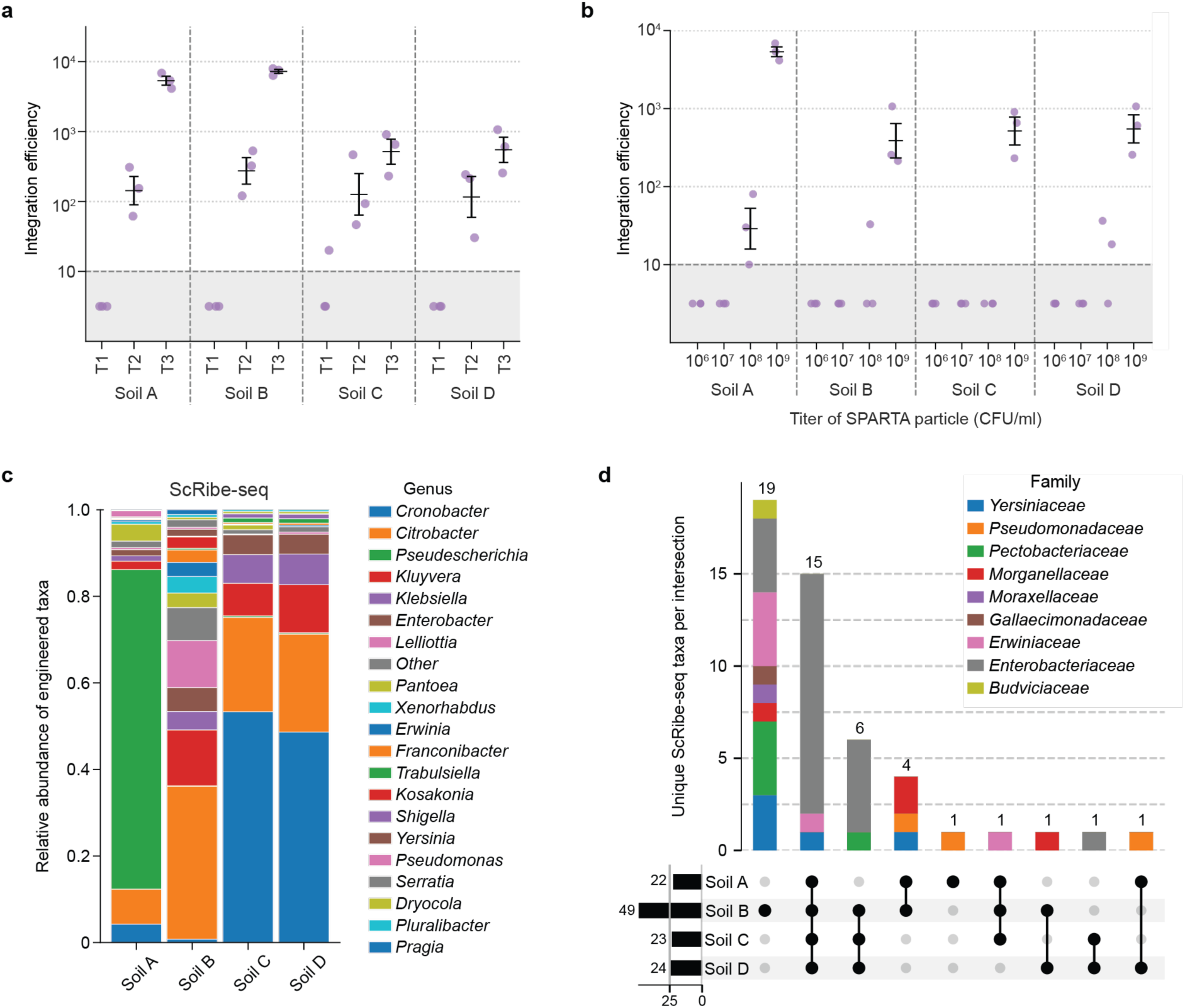
Detailed characterization of *in situ* soil SPARTA engineering. Integration efficiency of soil microbiomes mediated by SPARTA particles, shown as a function of **(a)** the number of treatments (T1–T3) and **(b)** SPARTA particle concentration (10^6^–10^9^ CFU/ml). **(c)** Genus-level composition of SPARTA engineered soil microbiome across soil samples assessed by ScRibe-seq. **(d)** UpSet plot showing the number of unique ScRibe-seq taxa (genus level) shared by each combination of soil samples. Bars are stacked by family-level composition within each intersection. All error bars indicate the mean ± standard error of the mean (SEM) from three experimental replicates (N = 3).

**Extended Data Figure 3.**
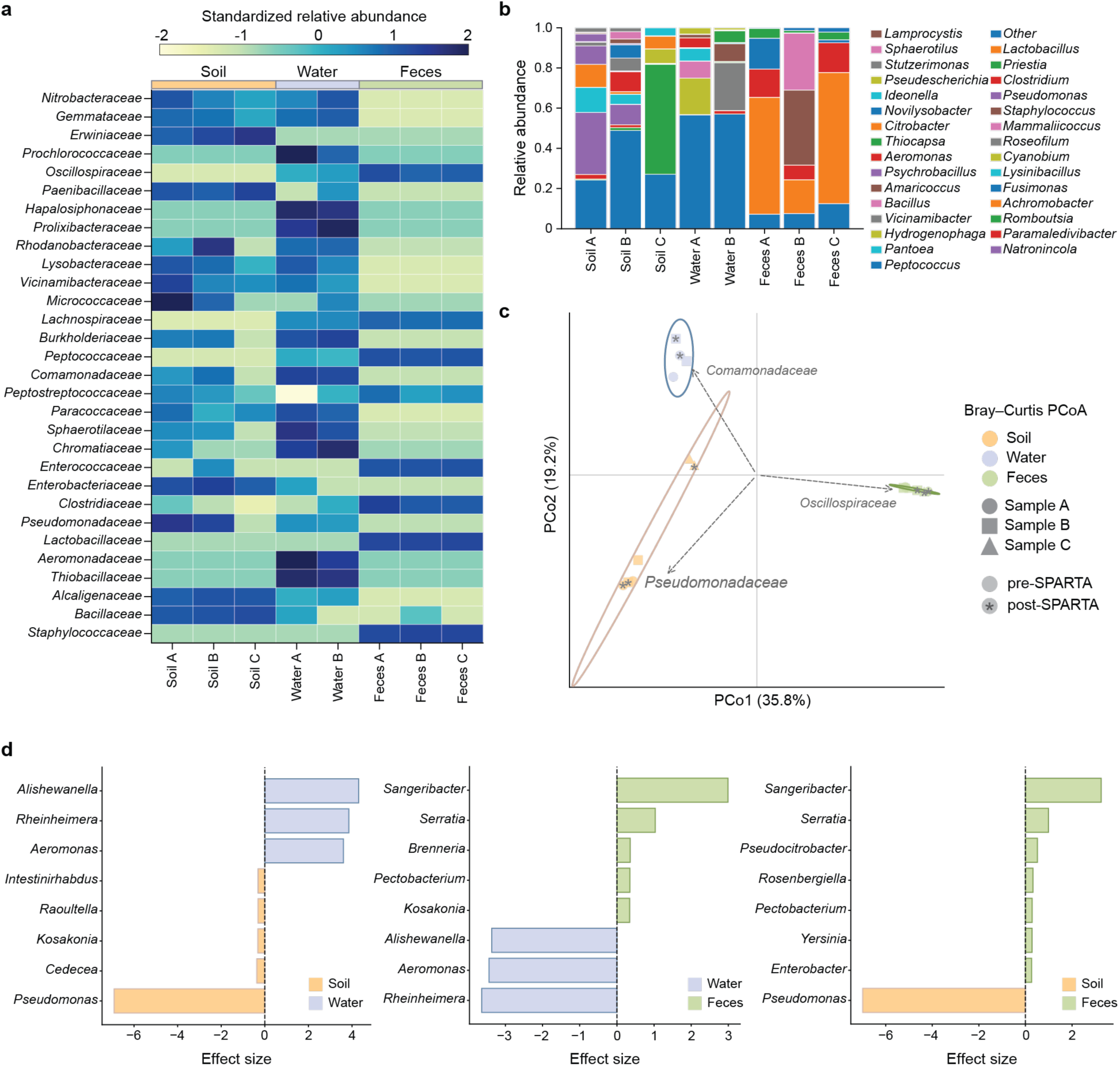
Characterization of SPARTA engineering with different sample types. **(a)** Standardized relative abundance of 16S rRNA taxa at the family level across sample types, showing distinct microbiome populations unique to each sample origin. **(b)** Genus-level microbiome composition in each sample quantified by 16S rRNA metagenomic sequencing. **(c)** Principal coordinate analysis (PCoA) by Bray-Curtis diversity of samples before and after SPARTA treatment, assessed by 16S rRNA sequencing (PERMANOVA, *P* = 0.0001, *R²* = 0.5375). Dashed arrows are representative correlation vectors for taxa with higher prevalence (>0.5). **(d)** ALDEX2 differential abundance analysis of ScRibe-seq taxa at the genus level across different samples engineered with SPARTA. The bars represent the median CLR effect size.

**Extended Data Figure 4.**
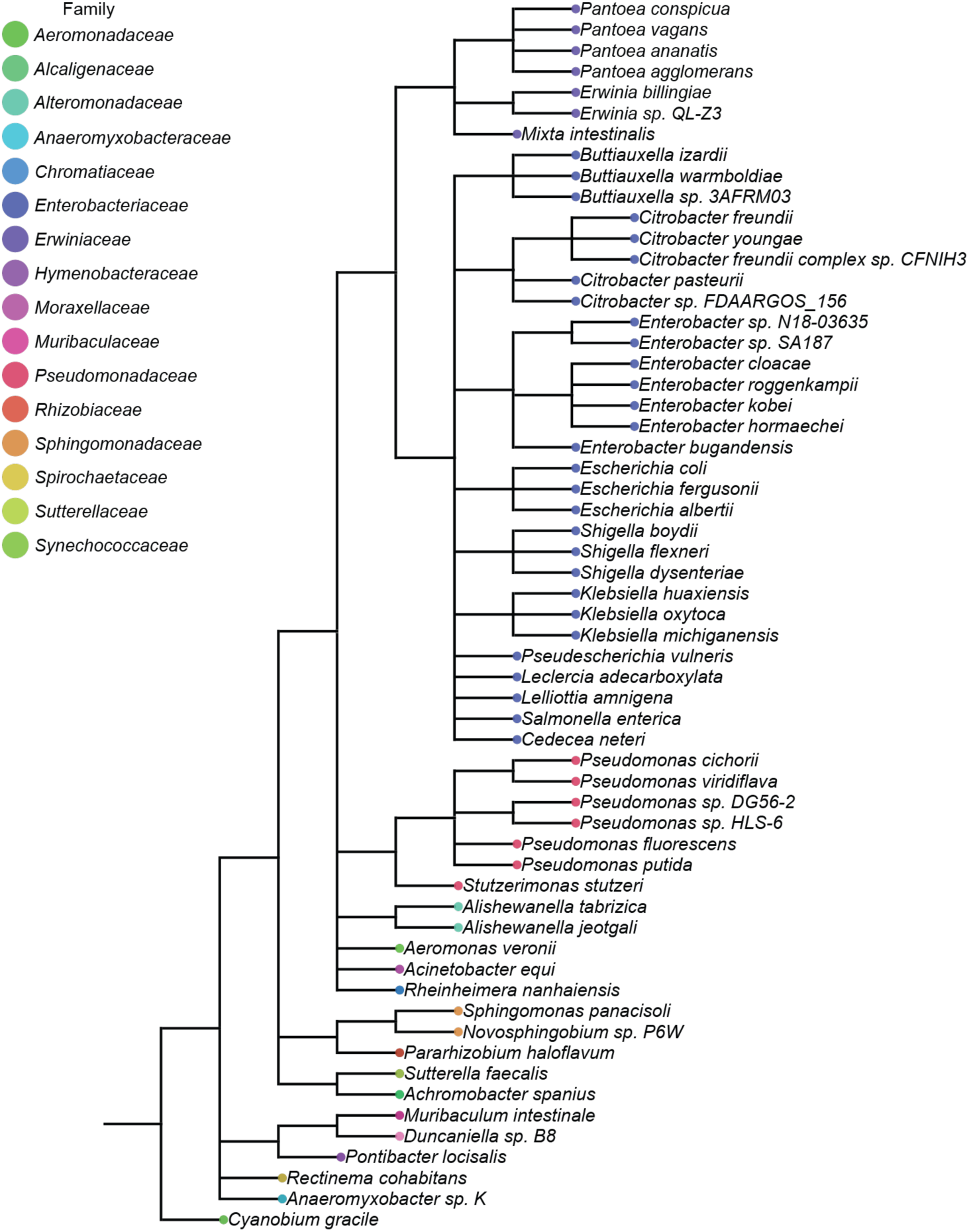
Phylogenetic tree of SPARTA engineered native microbiome species characterized by ScRibe-seq profiles using Emu pipeline. The color of the external nodes indicates higher-level taxonomy (family level).

**Extended Data Figure 5.**
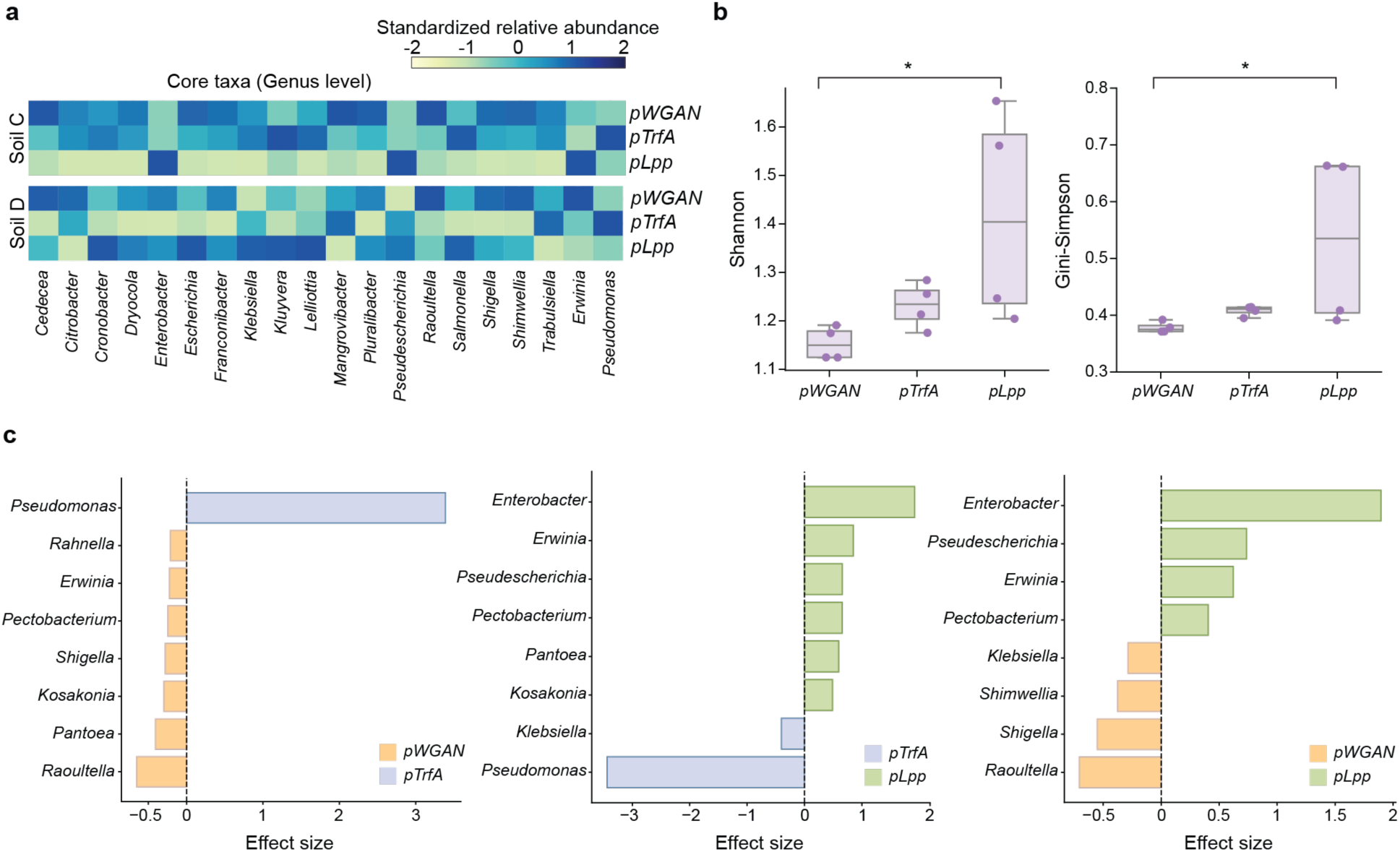
Description and validation of the genetic programmability of the SPARTA system. **(a)** Standardized relative abundance of representative ScRibe-seq taxa (genus level) identified across promoter variants for each soil sample. Representative taxa were defined as those with the highest mean relative abundance across all conditions within a given soil sample. **(b)** Shannon and Gini-Simpson diversity indices of ScRibe-seq taxa in soil samples treated with SPARTA variants (N = 4). The boxes span from the 25th to 75th percentiles (interquartile range), whiskers extend to 1.5 times the interquartile range and the center lines represent the median. **(c)** ALDEx2 differential abundance analysis of ScRibe-seq taxa at the genus level across different promoters of SPARTA. The bars represent the median CLR effect size.

**Extended Data Figure 6.**
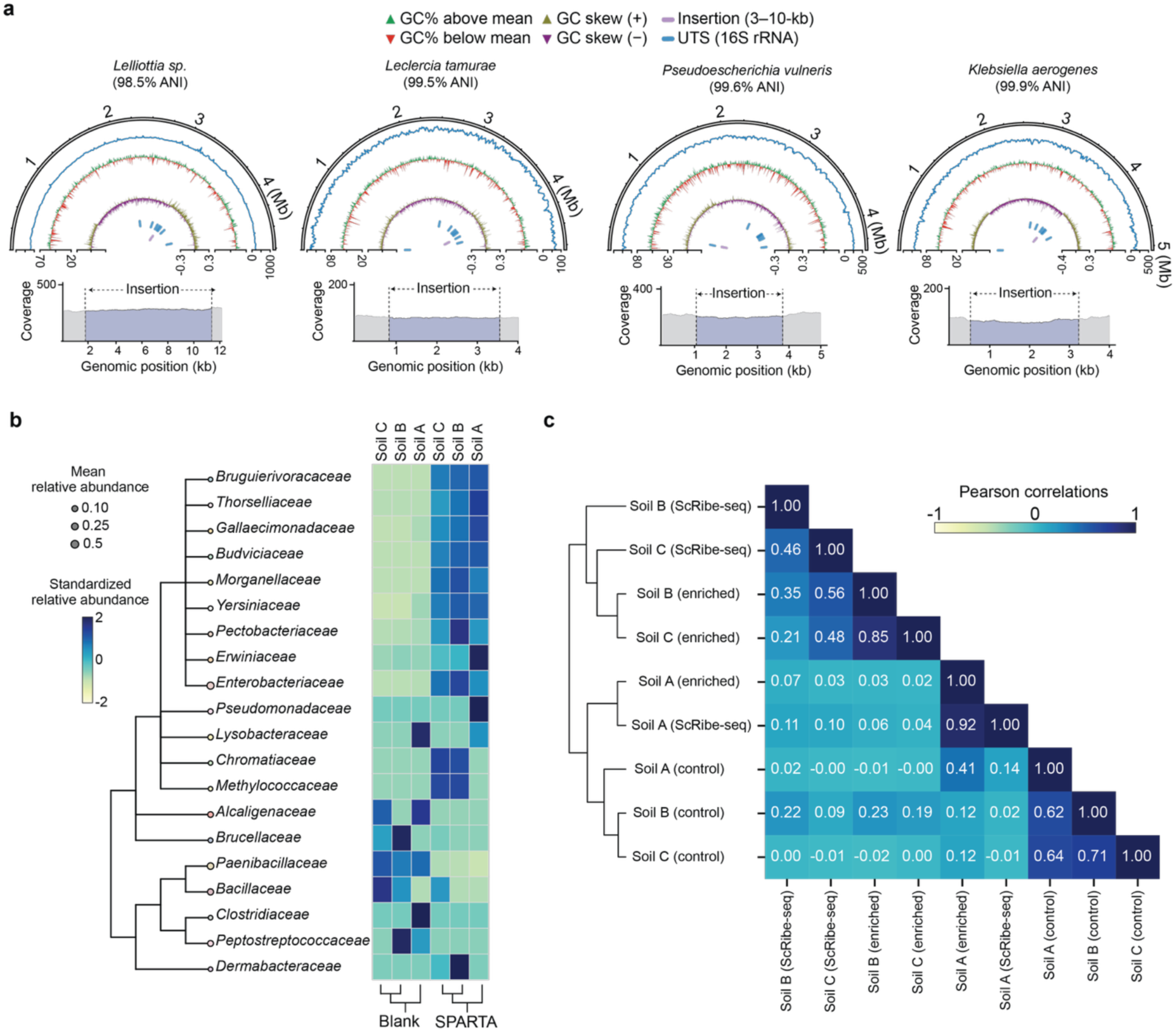
Isolation and selective enrichment of SPARTA engineered soil. **(a)** Whole genome sequencing validation of SPARTA-directed genome editing in isolated soil bacterial species. Semicircular plots present read coverage, GC content (%), and GC skew (from outer to inner rings) along the genomic position and detailed coverage plot of the insertion site is shown below. Blue and purple features indicate the target locus (16S rRNA) and the cargo insertion site, respectively, on the genome. ANI denotes average nucleotide identity estimated by sourmash. **(b)** Standardized relative abundance of microbiome taxa at the family level from soil cultures grown with antibiotics following blank or SPARTA treatment. Node size indicates the mean relative abundance of taxa across samples. **(c)** Correlation matrix comparing ScRibe-seq profiles of SPARTA treated soil samples with conventional 16S rRNA sequencing profiles of corresponding soil controls and antibiotic-enriched soil cultures.

**Extended Data Figure 7.**
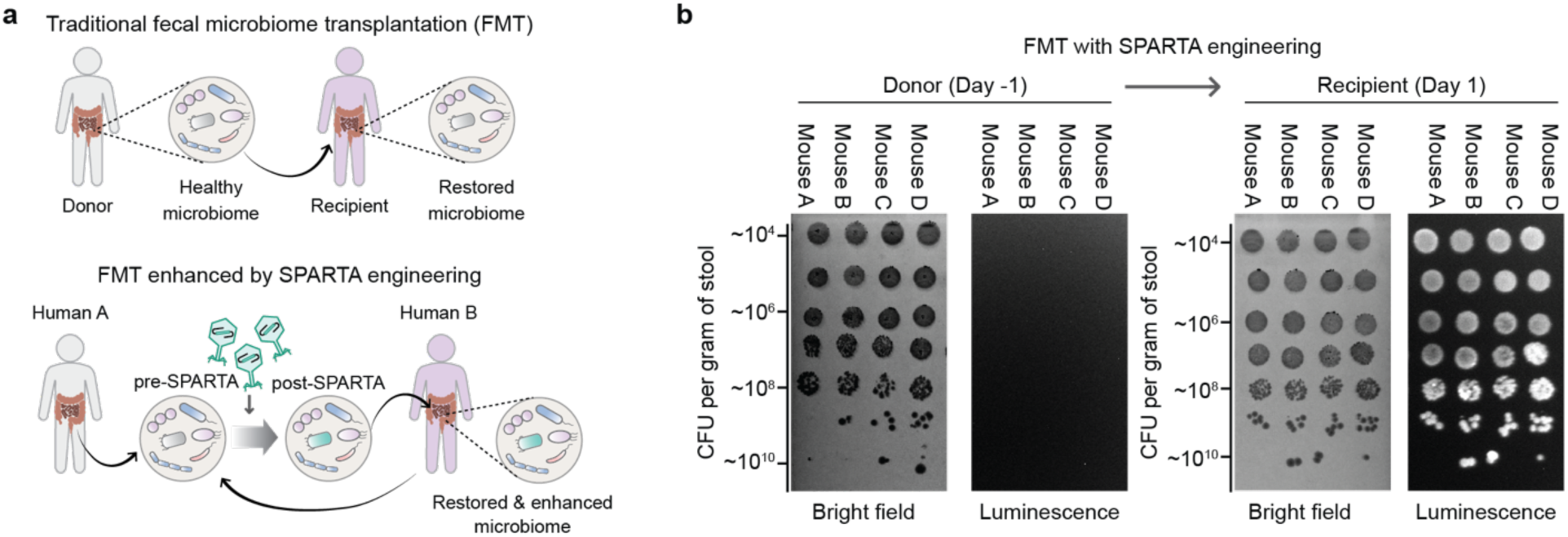
(a) Schematic of SPARTA engineering during the FMT process to enhance its therapeutic potential. **(b)** Representative bioluminescence phenotyping assay of transferred *E. coli* cells detected in stool samples from the donor (pre-FMT) and recipient (post-FMT).

**Extended Data Figure 8.**
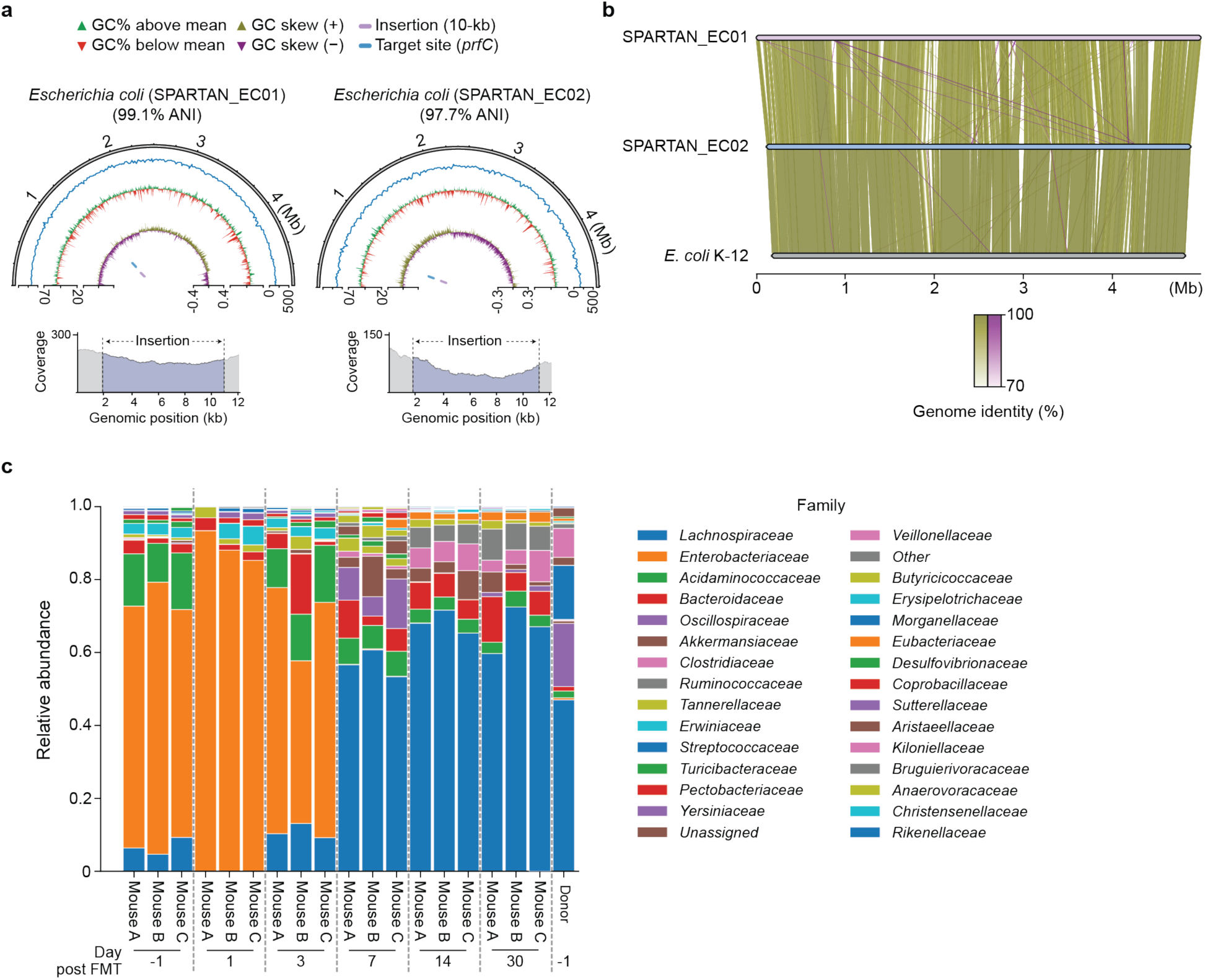
Detailed characterization of SPARTA engineered human fecal samples transplanted into recipient mice. **(a)** Whole genome sequencing of SPARTA engineered human native *E. coli* cells (e.g., SPARTAN_EC01–02) isolated from recipient mice stool samples following human-to-mouse fecal transplantation. Semicircular plots present read coverage, GC content (%), and GC skew (from outer to inner rings) along the genomic position and detailed coverage plot of the insertion site is shown below. Blue and purple features indicate the target locus (*prfC*) and the cargo insertion site, respectively. **(b)** Pairwise alignments between genomes of the two isolated human native *E. coli* strains and laboratory K-12 *E. coli*. Green links represent forward orientation alignment, and purple links indicate inverted orientation alignment. **(c)** Family-level gut microbiome composition of the human donor and recipient mice from one day before (Day-1) to 30 days after (Day 30) transplantation.

## Notes

### Competing Interest Statement

The authors have declared no competing interest.

